# Integrative transcriptomic identification of cellular senescence beyond marker limitations

**DOI:** 10.64898/2026.01.02.697374

**Authors:** Jing Lu, İsmail Güderer, Tayyaba Alvi, Mark Olenik, Handan Melike Dönertaş

## Abstract

Cellular senescence lacks a universal marker and varies across cell types, tissues, and stressors, complicating identification. Using SPiDER SA-β-gal labeled single-cell RNA-seq from regenerating mouse muscle, we found that curated gene sets show opposing enrichment patterns in experimentally defined senescent cells, suggesting apparent concordance in prior studies may reflect circular validation. Machine learning classifiers outperformed marker-centric approaches by capturing coordinated transcriptional features largely absent from differentially expressed genes. These features traced senescence progression, positioning senescent cells at late pseudotime with reduced transcriptional entropy. Ligand-receptor analysis identified IGF signaling as a directional axis of secondary senescence from senescent to non-senescent cells. When applied to bulk RNA-seq and an independent aging dataset, the classifier detected age-associated senescence patterns while the entropy-senescence relationship held across most cell types. These findings demonstrate that transcriptome-based classification provides a robust alternative to marker-centric readouts while enabling mechanistic hypothesis generation.

## Introduction

Cellular senescence, triggered by persistent DNA damage and other stress signals, including oxidative and oncogenic stress, is characterized by stable cell-cycle arrest coupled with sustained metabolic activity^1^. It contributes to organismal aging and multiple age-associated pathologies, including cancer, cardiovascular disease, and neurodegeneration^2^. Senescent cells can influence neighboring cells through paracrine signaling. This process, referred to as secondary senescence, involves the reinforcement or propagation of the senescence state via the senescence-associated secretory phenotype (SASP)^3,4^. Therapeutic efforts accordingly target either the selective elimination of senescent cells (senolytics) or the modulation of their pro-inflammatory secretome (senostatics or senomorphics)^5^.

The concept of cellular senescence traces back to the Hayflick limit^6^, which described the finite replicative capacity of human fibroblasts and later linked replicative exhaustion to telomere attrition^7^. Subsequent work has expanded the phenotype to include lysosomal expansion, chromatin remodeling, SASP induction, metabolic shifts, and characteristic morphological changes, in addition to durable proliferation arrest^8,9^. This breadth is biologically informative yet methodologically challenging: no single feature is sufficient or necessary across tissues, species, or inducing stimuli^10^. Accordingly, identification typically relies on multiple markers interpreted in context. Common readouts include senescence-associated β-galactosidase (SA-β-gal) activity and cell-cycle inhibitors (p16 and p21), alongside SASP components such as cytokines (e.g., IL6, IL8, CCL chemokines, TNF-α), growth factors (e.g., VEGF, HGF, TGF-β, IGFBPs), and proteases (e.g., MMPs, PAI-1)^11^. To structure evidence while acknowledging heterogeneity, Gil and colleagues proposed a three-tier framework^8^. This framework distinguishes primary markers (e.g., SA-β-gal, lipofuscin), validation markers (e.g., p16, p21, Ki67/BrdU negativity, Lamin B1), and characterization markers (e.g., SASP modules, DNA-damage response, subtype-specific features). Senescence heterogeneity is a central obstacle^12,13^. Senescence varies by cell type, tissue, species, inducing stressor (e.g., DNA damage, oncogenic signaling, organelle stress), and intervention (e.g., senolytics vs. senostatics), and it differs between in vitro and in vivo contexts. These sources of variation complicate reproducible classification and limit the portability of single-marker approaches across datasets^14^.

High-throughput technologies now offer new avenues to address these challenges. Single-cell RNA-seq (scRNA-seq), bulk transcriptomics, and high-parameter cytometry enable simultaneous measurement of many candidate markers at cellular resolution^15,16^. In this setting, curated gene sets can aggregate weak signals. For example, SenMayo and related resources such as GenAge and CellAge have shown utility relative to single markers, though their content and performance can differ across datasets^17^. However, despite these advantages, in practice, dataset-internal thresholding of enrichment scores can also introduce circularity when labels are derived from the very scores being evaluated.

To address these challenges, we integrate scRNA-seq with an extension of the traditional SA-β-gal assay. SPiDER SA-β-gal provides improved cell permeability, intracellular retention, rapid staining kinetics, and live-cell compatibility, allowing downstream single-cell sequencing^18^. Using SPiDER-based labels in the context of injury-induced senescence in mouse muscle regeneration^15^, we benchmark individual markers against curated senescence/aging gene sets (SenMayo, GenAge, CellAge) and evaluate a composite ratio that seeks to combine complementary signal directions. We then train machine-learning classifiers for senescence by leveraging the full transcriptome of single cells. We examine which transcriptomic features the models consistently prioritize and whether these signals track senescence progression. This design aims to improve practical classification while retaining biological interpretability grounded in experimentally supported labels, and to identify pathways implicated in cellular senescence and secondary senescence transduction. In addition, we applied our classifier alongside other published machine learning (ML) approaches to the Tabula Muris Senis aging dataset, where ENet detected age-associated senescence increases in several cell types and identified a low-entropy transcriptional state in predicted senescent cells across most cell-types examined.

## Results

### Data description

We analyze single-cell RNA-seq from regenerating mouse skeletal muscle (tibialis anterior) at 3 days post cardiotoxin (CTX) injury in young adults (3-6 months)^15^. Senescent and non-senescent cells were prospectively isolated by FACS using the SPiDER-β-gal probe, sorting both CD45⁺ and CD45⁻ compartments, and then merging fractions for profiling. The resulting atlas comprises 21,239 cells resolved into major lineages: myeloid cells (MCs), fibro-adipogenic progenitors (FAPs), and satellite cells (SCs). This design provides paired SPiDER⁺/SPiDER⁻ subsets within each cell-type (N_FAP:Sen_= 6,534, N_FAP:NSen_= 3,307, N_MC:Sen_= 6,796, N_MC:NSen_= 1,897, N_SC:Sen_= 576, N_SC:NSen_= 2,129), enabling experimentally labelled classification of senescence states. Importantly, the dataset includes two biological replicates, each preprocessed and sequenced separately. In our analysis, we use both replicates, which allows us to test reproducibility and hold-out testing in machine learning models.

### Curated senescence marker sets outperform single markers

We first evaluated classical single-gene markers using SPiDER SA-β-gal as the experimental label across both biological replicates jointly. *Cdkn2a, Cdkn1a, and Tgfb1* showed minimal separation between SPiDER-negative and SPiDER-positive cells (Average log_2_ fold-change for *Cdkn1a* = -0.106, *Cdkn2a* = -0.083, *Tgfb1* = 0.211), with *Cdkn2a* expressed at low levels and in only a small subset of cells **(Fig. 1a)**. This observation is consistent with prior reports of inconsistent performance of single markers for senescence detection in transcriptomic data^17,19,20^.

**Figure 1:**
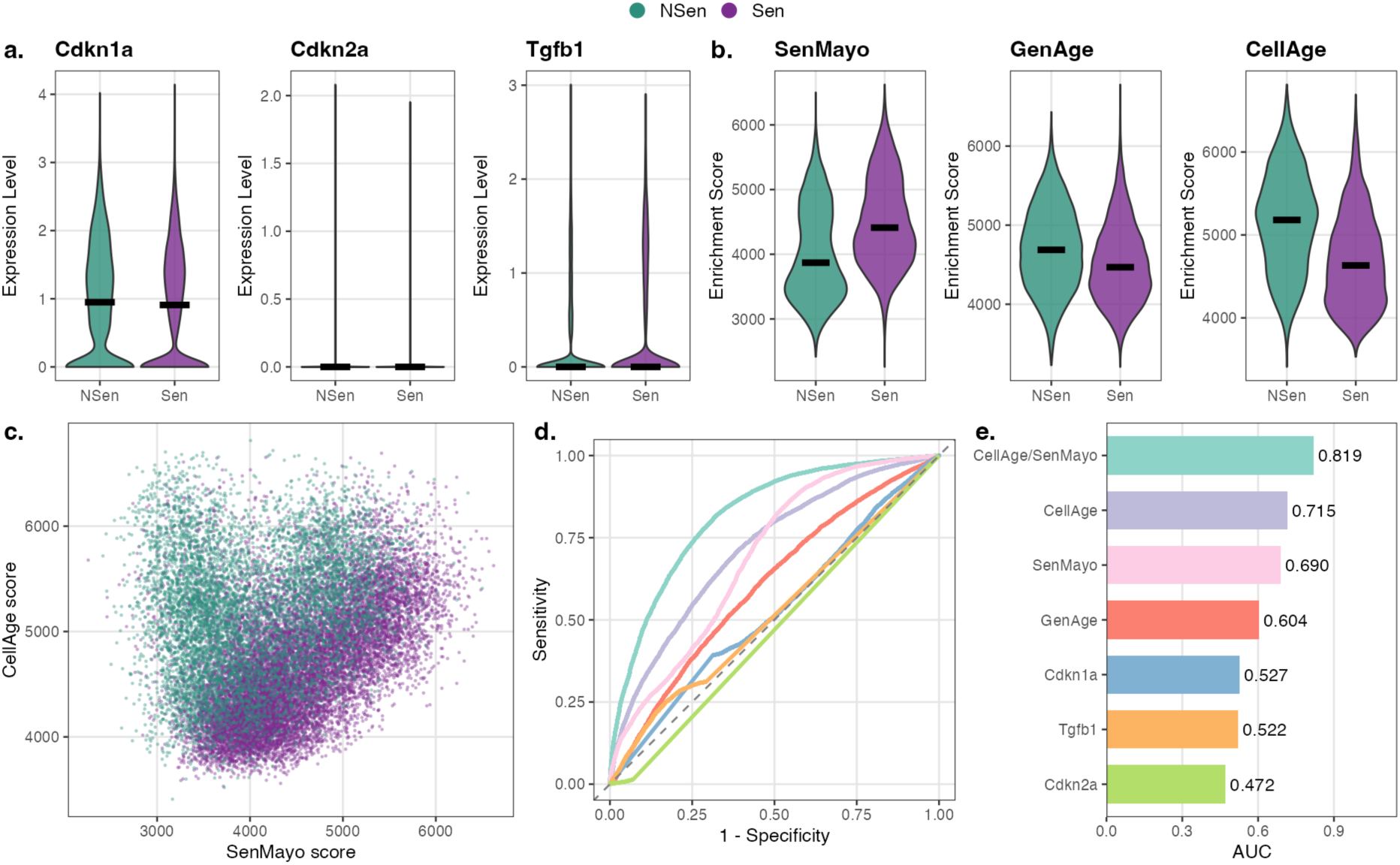
Curated senescence marker sets outperform single markers across both biological replicates. **a)** Violin plots showing expression of classical senescence markers Cdkn1a, Cdkn2a, and Tgfb1 in SPiDER-negative (NSen) and SPiDER-positive (Sen) cells. Cdkn2a was expressed at low levels in only a small subset of cells, while Cdkn1a and Tgfb1 showed minimal separation between senescent and non-senescent populations. **b)** Per-cell enrichment scores for curated senescence/aging gene sets: SenMayo (mouse), GenAge (mouse), and CellAge, stratified by SPiDER-defined senescence status. SenMayo was enriched in SPiDER-positive cells, whereas GenAge and CellAge showed enrichment in SPiDER-negative cells. **c)** Scatter plot comparing SenMayo and CellAge enrichment scores across single cells. Unlike prior enrichment-based labeling approaches, no co-enrichment was observed when using experimental SPiDER labels. **d)** Receiver operating characteristic (ROC) curves for enrichment scores of marker sets and expression of individual markers. Colors correspond to predictors shown in panel e. **e)** Area under the ROC curve (AUC) values for each predictor. Single markers had little discriminatory power (AUC ≈ 0.5), whereas curated gene sets showed higher accuracy (CellAge: 0.715; SenMayo: 0.690; GenAge: 0.604). A composite ratio of CellAge to SenMayo enrichment achieved the best performance (AUC = 0.819).

To test whether aggregating signals improves discrimination, we computed per-cell enrichment scores for three curated aging and senescence gene sets (**Supplementary Table S1**), SenMayo (mouse)^17^, GenAge (mouse), and CellAge^21^ **(Fig. 1b)**. The SenMayo gene set was curated from 15 key studies by screening 1,656 studies to identify genes (n=118, mouse) consistently enriched in senescent and SASP-secreting cells, with experimental validation in at least human or mouse cells. GenAge is a curated database of genes (n=136) associated with aging and longevity, compiled through an extensive literature review across model organisms and humans. CellAge focuses on genes (n=866) that regulate cellular senescence, identified through experimental evidence and manual curation of studies involving human cell lines. The three resources overlap only partially, with CellAge substantially larger than the others, indicating distinct curation scopes **(Extended Data Fig. 1a)**. Using SPiDER-defined labels, we observed higher SenMayo scores in senescent cells, whereas GenAge and CellAge scores were lower relative to non-senescent cells **(Fig. 1b**, Average log₂ fold-change: SenMayo = 0.165, GenAge = -0.057, CellAge = -0.136; Wilcoxon rank-sum test p < 2.2×10⁻¹⁶). This divergence suggests that the sets capture different facets of senescence. The SenMayo set primarily reflects the SASP and is enriched in SPiDER-identified senescent cells, whereas GenAge and CellAge sets, which contained both longevity-related genes (with inhibited senescence effects) and aging-related genes (with induced senescence effects), show overall the opposite pattern. Because earlier studies often defined “senescent” cells by dataset-internal SenMayo thresholds, we reproduced that approach: labeling cells by the top decile of SenMayo enrichment recapitulated the originally reported co-enrichment of GenAge and CellAge with SenMayo **(Extended Data Fig. 1b-c)**^17^. In contrast, when using experimental SPiDER labels, GenAge/CellAge did not co-enrich with SenMayo **(Fig. 1c)**. This discrepancy highlights the risk of circularity when enrichment scores serve as both labels and evaluation metrics, and underscores the need for experimental ground truth.

To quantify the classification performance, we computed receiver operating characteristic (ROC) curves for enrichment scores and individual marker expression levels **(Fig. 1d-e)**. Single markers showed no classification power (AUC ∼ 0.5), while marker sets demonstrated higher performance, with AUC values of 0.715, 0.690, and 0.604 for CellAge, SenMayo, and GenAge, respectively. Interestingly, most separations arose from FAPs and MCs, while SCs showed weaker SenMayo enrichment irrespective of SPiDER status. The average Euclidean centroid distance in the enrichment feature space between senescent and non-senescent cells was 527 in FAPs, 584 in MCs, and 213 in SCs, highlighting the markedly reduced separation in SCs **(Extended Data Fig. 1d)**. We next assessed whether the lack of concordance between SenMayo and CellAge scores in indicating senescence state was specific to the dataset we analysed. We include an independent WI-38 fibroblast dataset (GSE226225)^22^ to validate the enrichment relationship between SenMayo and CellAge. The dataset (GSE226225) spanning control (proliferating cells or ETO day0), replicative senescence (RS), irradiation-induced senescence (IR), and etoposide-induced senescence (ETO), senescent cells distributed along a CellAge–SenMayo diagonal rather than forming a SenMayo-high cluster alone **(Extended Data Fig. 1e–h)**. Motivated by the complementary behavior of the gene sets, we built a composite baseline by using the ratio of CellAge to SenMayo enrichment score, which challenges existing approaches that define senescence based on concordance across multiple marker gene sets^23^. This ratio achieved the highest baseline performance (AUC = 0.819), exceeding any single set and all single-marker baselines **(Fig. 1d-e)**. Taken together, these results indicate that curated marker sets outperform single genes for SPiDER-labelled classification, and that combining sets can reconcile partially conflicting aspects of the senescence signal. However, the limitations of thresholded enrichment-based labeling, particularly the risk of circular evaluation, motivate the development of machine learning approaches trained on experimental labels.

### Machine learning models outperform standardized baselines and reveal biologically structured features

To test whether machine learning (ML) classifiers outperform standardized baselines, we trained three models (elastic net (ENet), random forest (RF), and support vector machine (SVM)) on one sample (Sample 2, n = 11,612 cells; selected for training due to larger size) and evaluated performance on an independent held-out sample (Sample 1, n = 9,627 cells). For direct comparison, all AUC values reported in Fig 1d-e, were re-computed on the same test sample (**Fig. 2a**).

**Figure 2:**
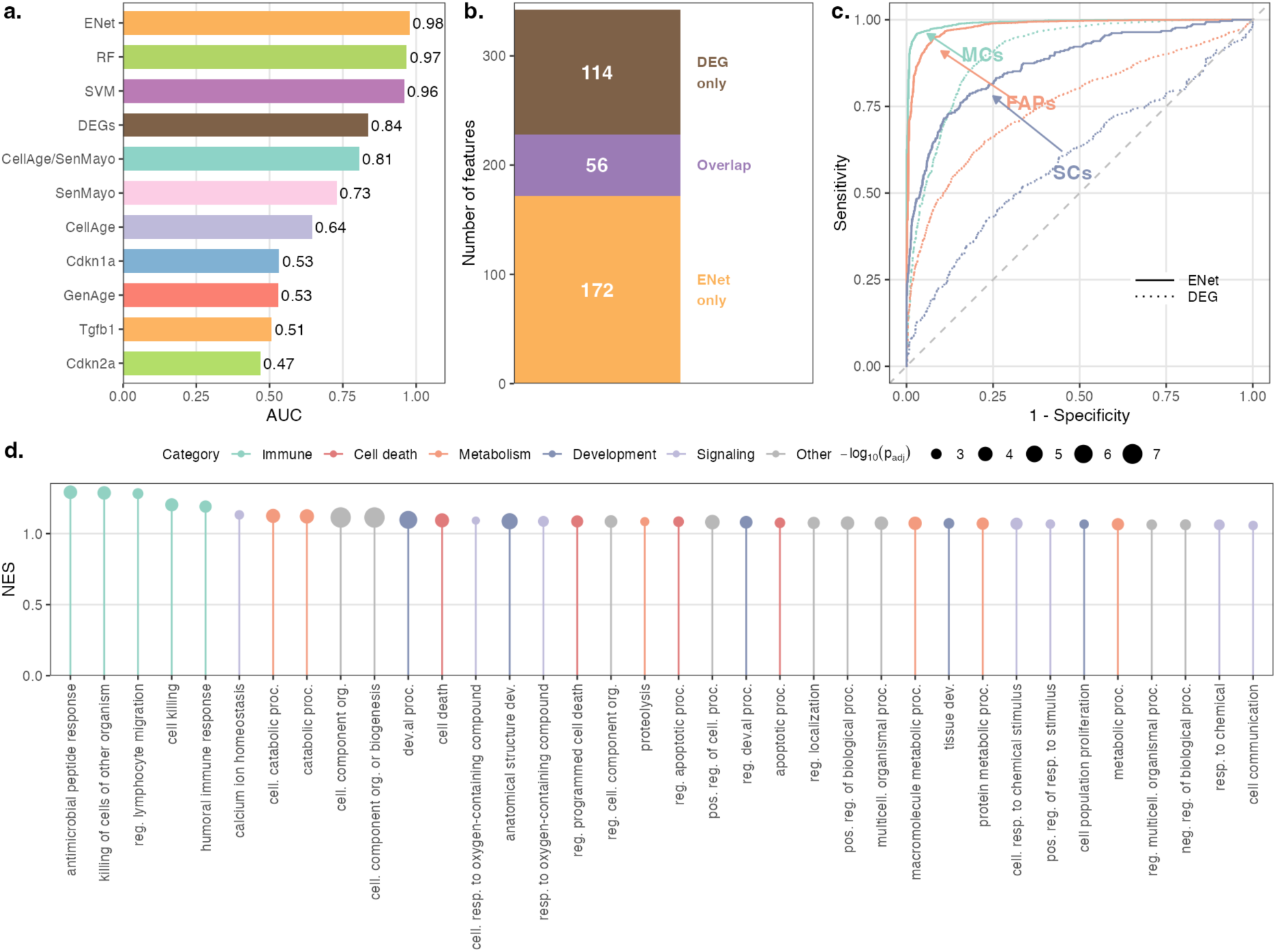
Machine learning models outperform standardized baselines and reveal biologically structured features. **(a)** Classification performance (AUC) of machine learning models (ENet, RF, SVM) compared to DEG-based predictors and curated gene sets in the test sample. All ML models achieve AUC > 0.95, outperforming standardized baselines (DEG: 0.84; CellAge/SenMayo: 0.81). **(b)** Composition of ENet-important features (n = 228) relative to significant DEGs (n = 170). Only 56 features overlap; 172 ENet-important features were not identified by differential expression analysis. **(c)** ROC curves comparing ENet (solid lines) and DEG-based (dashed lines) classification across cell types in the test sample: myeloid cells (MCs, teal), fibro/adipogenic progenitors (FAPs, coral), and satellite cells (SCs, slate blue). ENet improves prediction in all cell types, with the largest relative gain in SCs (AUC: ENet = 0.874 vs DEG = 0.599). **(d)** Gene ontology enrichment analysis of ENet-important features (Biological Process). Terms ranked by normalized enrichment score (NES), colored by manual annotation of biological categories, with point size proportional to significance (−log₁₀ Padj).

Consistent with Fig. 1, single-gene markers showed no discriminatory power for senescence classification (AUC range: 0.47 to 0.53; **Fig. 2a**). As standardized baselines, a DEG set defined on the training data (Bonferroni-adjusted P < 0.05; absolute log₂ fold-change above the elbow threshold, n = 170 genes; **Extended Data Fig. 2a**) achieved AUC = 0.84, and the CellAge/SenMayo composite score achieved AUC = 0.81. All three ML models exceeded these baselines with AUC_ENet_ = 0.98, AUC_RF_ = 0.97, and AUC_SVM_ = 0.96 **(Fig. 2a)**. Feature importance profiles differed across models and diverged from univariate rankings based on AUC or log₂ fold-change (**Extended Data Fig. 2d; Supplementary Table S2**). Given its high performance and the inherent interpretability of linear models, we focused downstream analyses on ENet.

We defined ENet-important features as those with absolute coefficients exceeding the elbow point threshold (n = 228 features; **Extended Data Fig. 2b; Supplementary Table S2**). Among these 228 features, only 56 overlapped with significant DEGs (n = 170), indicating that the model captures information beyond conventional differential expression (**Fig. 2b**). Within the important feature set, pairwise expression correlations were significantly stronger than correlations within the non-important set or correlations between important and non-important features (P < 2.2 × 10⁻¹⁶ for all comparisons, Wilcoxon rank-sum test; **Extended Data Fig. 2c**). This pattern suggests coordinated expression among prioritized genes, consistent with structured, potentially functional modules.

Classification performance was high across all three cell types, with ENet achieving AUC = 0.991 for MCs and AUC = 0.981 for FAPs (**Fig. 2c**). SCs presented the greatest classification challenge, consistent with their smaller sample size (n = 576 senescent, n = 2,129 non-senescent) and class imbalance. Indeed, DEG-based classification performed near chance for SCs (AUC = 0.599). The ENet model substantially improved SC classification (AUC = 0.874), representing the largest relative gain across cell types. A potential concern with ML classifiers is that cell-type identity might confound senescence-state classification. However, if this were the case, stratifying the analysis by cell type should lead to a drop in performance, which we do not observe. Instead, cell-type-stratified AUCs remained close to the combined AUC, indicating that the model captures senescence state within each lineage. To further assess potential cell-type confounding, specifically, whether our ML features reflect cell-type identity rather than senescence, we tested whether ENet-important features were enriched for cell-type markers after filtering out all senescence-related genes. FAP markers showed significant enrichment (OR = 2.08, P = 0.017), MC markers showed depletion (OR = 0.81, P = 0.83), and SC markers showed non-significant enrichment (OR = 1.95, P = 0.24; **Extended Data Fig. 2e,f**). This enrichment pattern did not correspond to classification performance: FAPs and MCs achieved comparable AUCs despite opposite marker enrichment patterns, and SCs showed the largest performance gain despite lacking significant marker enrichment. Gene ontology analysis of ENet-important features identified established senescence processes, including immune and inflammatory processes, cell death pathways, stress responses, metabolic and catabolic programs, and tissue remodeling (**Fig. 2d**).

To assess whether a compact classifier preserves performance, we applied recursive feature elimination (RFE) to the ENet model using repeated 10-fold cross-validation on training data. Model size was selected by an elbow criterion on accuracy versus feature count. This procedure identified an optimal 100-feature model that achieved 96% accuracy while reducing the feature set by 56% as compared to the 228 feature model (**Extended Data Fig. 2g**). Across resampling schemes, RFE identified 147 recurrent features, of which 139 (95%) overlapped with the top-ranked features from the initial ENet classifier (**Extended Data Fig. 2h**). This convergence supports the reproducibility of ENet-prioritized features across resampling schemes, indicating that they capture genuine biological signals rather than noise.

Together, these results demonstrate that ML models, including a simple regularized ENet, surpass standardized baselines on independent held-out data. The models recover structured multi-gene signals that univariate approaches miss and deliver substantial improvements in the most challenging cell type.

### ML-prioritized features reconstruct senescence-aligned pseudotime trajectories and reveal late-stage heterogeneity

To test whether model-selected features capture progression rather than classification alone, we constructed pseudotime trajectories using ENet-important features (n = 228) from integrated data across both biological replicates, applying Monocle2 over a UMAP embedding. With transcriptional entropy measuring the diversity and disorder of gene expression within a cell, we selected the trajectory root using the maximum entropy criterion with TSCAN, an unsupervised approach motivated by higher transcriptional entropy in less-committed states^21^ (**Extended Data Fig. 3a**). Across all three cell types, the ENet-important features traced late-stage branches enriched for SPiDER-positive cells (**Fig. 3a–c**). Senescent cells occupied significantly higher pseudotime than non-senescent cells within each cell type (Wilcoxon rank-sum test, P < 2.2 × 10⁻¹⁶ for all comparisons; **Fig. 3d–f**).

**Figure 3:**
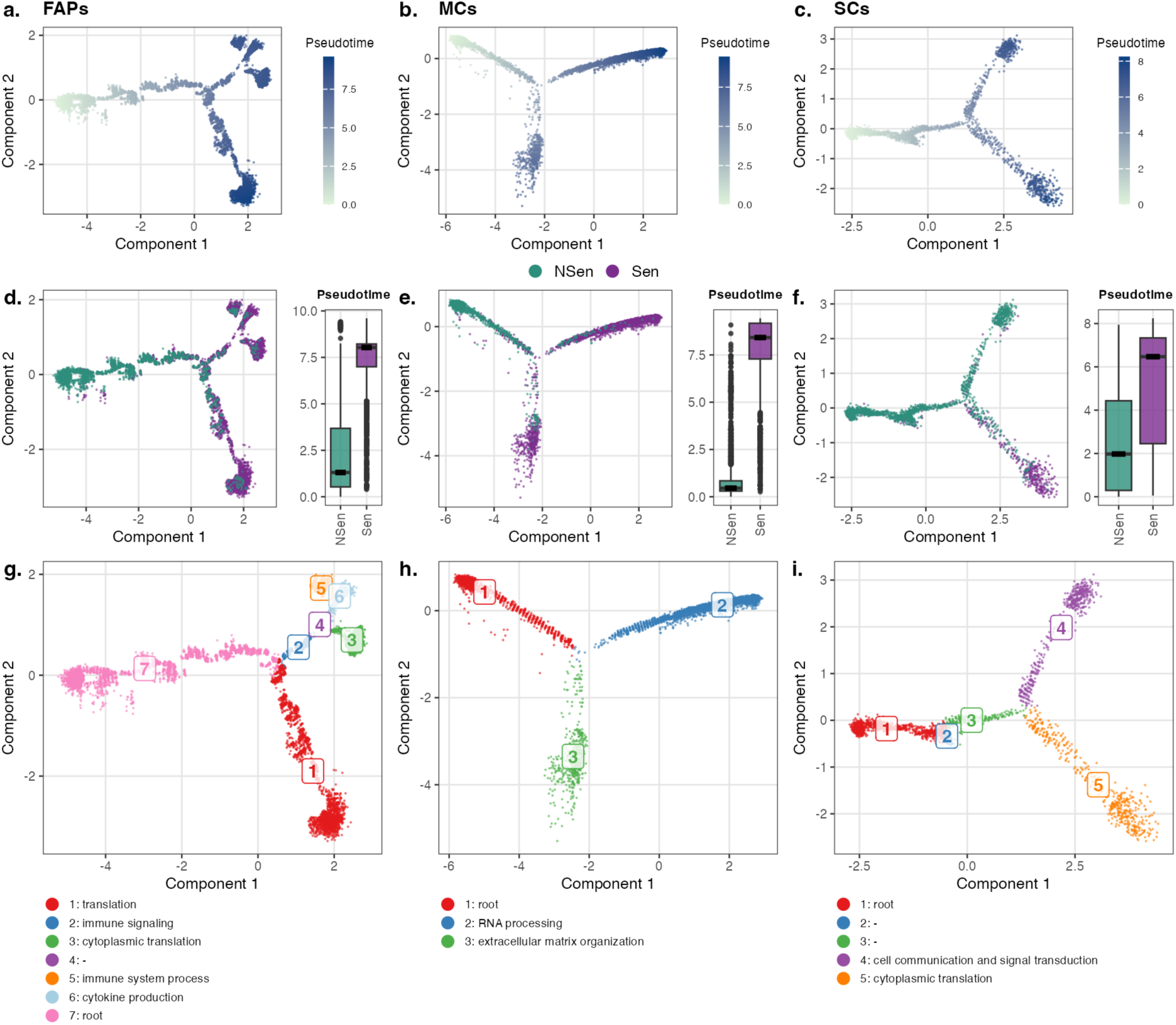
ML-prioritized features reconstruct senescence-aligned pseudotime trajectories and reveal late-stage heterogeneity. **(a-c)** Pseudotime trajectories constructed from ENet-important features (top 228) using Monocle2 over UMAP embeddings for FAPs (a), MCs (b), and SCs (c). Color gradient indicates pseudotime values (green = early, blue = late); each cell type has its own pseudotime scale. **(d-f)** Same trajectories colored by senescence state (teal = NSen, purple = Sen) for FAPs (d), MCs (e), and SCs (f). Adjacent boxplots compare pseudotime distribution between senescent and non-senescent cells; black crossbars indicate median values. Senescent cells occupy significantly higher pseudotime in all cell types (Wilcoxon rank-sum test, p < 2.2×10⁻¹⁶ for all comparisons). **(g-i)** Trajectory state annotations with GO enrichment for late branches compared to root state. (g) In FAPs (7 states), late states are enriched for immune signaling (state 2), translation (state 1), immune system process (state 5), cytokine production (state 6), and cytoplasmic translation (state 3). (h) In MCs (3 states), late states show enrichment for RNA processing (state 2) and extracellular matrix organization (state 3). (i) In SCs (5 states), the senescent branch (state 5) is enriched for cytoplasmic translation, while a non-senescent advanced branch (state 4) is enriched for cell communication and signal transduction.

In all three cell types, the ENet-important feature set outperformed less-important feature sets in recovering these late, senescence-enriched branches (**Extended Data Fig. 3c**). In MCs and FAPs, ENet-important feature performance was comparable to variable genes (**Extended Data Fig. 3c**). However, in SCs, ENet-important features substantially outperformed variable genes (AUC = 0.78 vs 0.66). These comparisons indicate that the ML-selected features carry information relevant to senescence progression beyond mere endpoint classification.

Pseudotime derived from ENet-important features correlated positively with the ENet classification score across cell types (FAPs: R = 0.85, p < 2.2 × 10⁻¹⁶; MCs: R = 0.93, p < 2.2 × 10⁻¹⁶; SCs: R = 0.42, p < 2.2 × 10⁻¹⁶; Extended Data Fig. 3d), yet the two quantities are distributed differently. Pseudotime spanned a continuous range, whereas senescence scores were more bimodal around the decision boundary. This divergence is expected because pseudotime encodes progression along a trajectory, whereas the ENet outputs a probability of class membership^24,25^.

Late-stage branches in the trajectory, which were predominantly enriched with senescent cells, further highlighted the diversity of senescent cells in FAPs and MCs (**Fig. 3d–e**), while showing reduced heterogeneity within senescent SCs (**Fig. 3f**). We compared each late senescent state with the root branch within each cell type to characterize functional differences between different states (**Supplementary Table S3**). In FAPs, one late state was enriched for immune response and signal-transduction pathways (state 2), while another was enriched for translation and peptide biosynthesis (state 1). A further split yielded sub-branches annotated by immune system process (state 5), cytokine production (state 6), and cytoplasmic translation (state 3; **Fig. 3g**). In MCs, late states were enriched for RNA processing and for extracellular matrix organization (**Fig. 3h**). In SCs, the senescent branch (state 5) showed enrichment for cytoplasmic translation, muscle system processes, and RNA processing, whereas a non-senescent advanced branch (state 4) was enriched for cell communication and signal transduction (**Fig. 3i**).

These annotations are consistent with context-dependent senescence programs^26^. We observed decreased transcriptional entropy in senescent cells compared with non-senescent cells. This pattern differs from aging-associated identity loss reported in some contexts^27,28^, though the relationship between acute senescence induction and organismal aging remains complex. This pattern was consistent across all three cell types in single-cell RNA-seq data (FAPs and MCs: p < 2 × 10⁻¹⁶; SCs: p = 6.3 × 10⁻⁴; Wilcoxon rank-sum test; **Extended Data Fig. 3b**) and in bulk RNA-seq data (FAPs: p = 6.3 × 10⁻⁵; MCs: p = 5.6 × 10⁻⁴; SCs: p = 1.7 × 10⁻⁴; Kruskal–Wallis test; **Extended Data Fig. 3e**). Notably, non-senescent cells following injury exhibited elevated entropy relative to non-injured contralateral controls. This pattern is consistent with tissue regeneration processes, during which cells may adopt less constrained transcriptional states^29–32^.

Together, these results indicate that ML-prioritized features recover a senescence-aligned trajectory that recapitulates experimental labels without using them for supervision. The analyses further highlight late-stage heterogeneity across lineages and reveal entropy dynamics that distinguish senescence from regeneration-associated transcriptional changes.

### Senescence-associated cell-cell interactions and IGF signaling in senescence transduction

To study secondary senescence, we performed gene set enrichment analysis (GSEA) on the ENet-prioritized feature set to nominate interaction-relevant pathway families (see Methods). We then inferred ligand-receptor (LR) communication with CellChat on the SPiDER-labelled single-cell data, stratifying emitters and receivers by senescence state.

Of the ten pathway families identified by GSEA, seven (SPP1, ApoE, FN1, PSAP, IGF, IGFBP, and CCL) showed significant LR communications by CellChat across senescence states and cell types (**Fig. 4a**). Several (e.g., SPP1^33^, ApoE^34^, IGF^35,36^, IGFBP^3,37^, and CCL^38^) have been previously linked to senescence and SASP. Some interaction pathways, including CCL, SPP1, and ApoE, were consistently identified through both traditional differential gene expression (DEG) analysis and ENet modeling (**Fig. 4b)**. However, other biologically relevant pathways (FN1, IGF, IGFBP, and SEMA3) emerged only through ENet feature prioritization. This highlights the added value of machine learning approaches, which uncover context-dependent signaling patterns by weighting features in ways that differ from conventional statistical methods, offering complementary insights into cell-cell communication that may be overlooked by DEG-based strategies alone. We examined results from other machine learning methods and found that IGF and FN1 signaling interactions were enriched among random forest features (**Extended Data Fig. 4a**) but were not prioritized by the SVM model (**Extended Data Fig. 4b**). IGF, SPP1, and SEMA3 ligands were predominantly expressed by senescent cells, with minimal expression in non-senescent populations (**Fig. 4c**, **Extended Data Fig. 4c**). Among these, SPP1 exhibited high inter-pseudostate variability with no consistent trajectory pattern across samples, whereas SEMA3 showed lower variability but inconsistent trajectories between biological replicates (**Extended Data Fig. 4d**). In contrast, inferred IGF interaction probability exhibited a reproducible trajectory across both biological replicates, increasing along pseudotime, peaking at pseudostates 6 to 7, and subsequently declining (**Fig. 4d**). Based on ligand-receptor expression patterns, senescent FAPs and MCs were predicted as the predominant IGF signal emitters, with SCs as primary receivers in both biological replicates (**Fig. 4c**). The *Igf1-Igf1r* ligand-receptor pair emerged as the principal contributor to IGF signaling-mediated cell-cell interactions across cell types and senescence states. Consistently, *Igf1* expression mirrored LR probability along pseudotime, whereas *Igf1r* expression peaked earlier than *Igf1* (**Fig. 4e**). *Igf1* expression was elevated in senescent FAPs and MCs (predicted emitters), whereas *Igf1r* was upregulated in SCs (predicted receivers), consistent with directional IGF signaling from senescent cells to satellite cells (**Fig. 4f**). Consistent with the antagonistic role of IGFBPs in modulating IGF bioavailability, IGF communication probability increased when IGFBPs were computationally masked, particularly for SCs as receivers (**Extended Data Fig. 4f**). This pattern supports IGFBP-mediated sequestration of IGF ligands.

**Figure 4:**
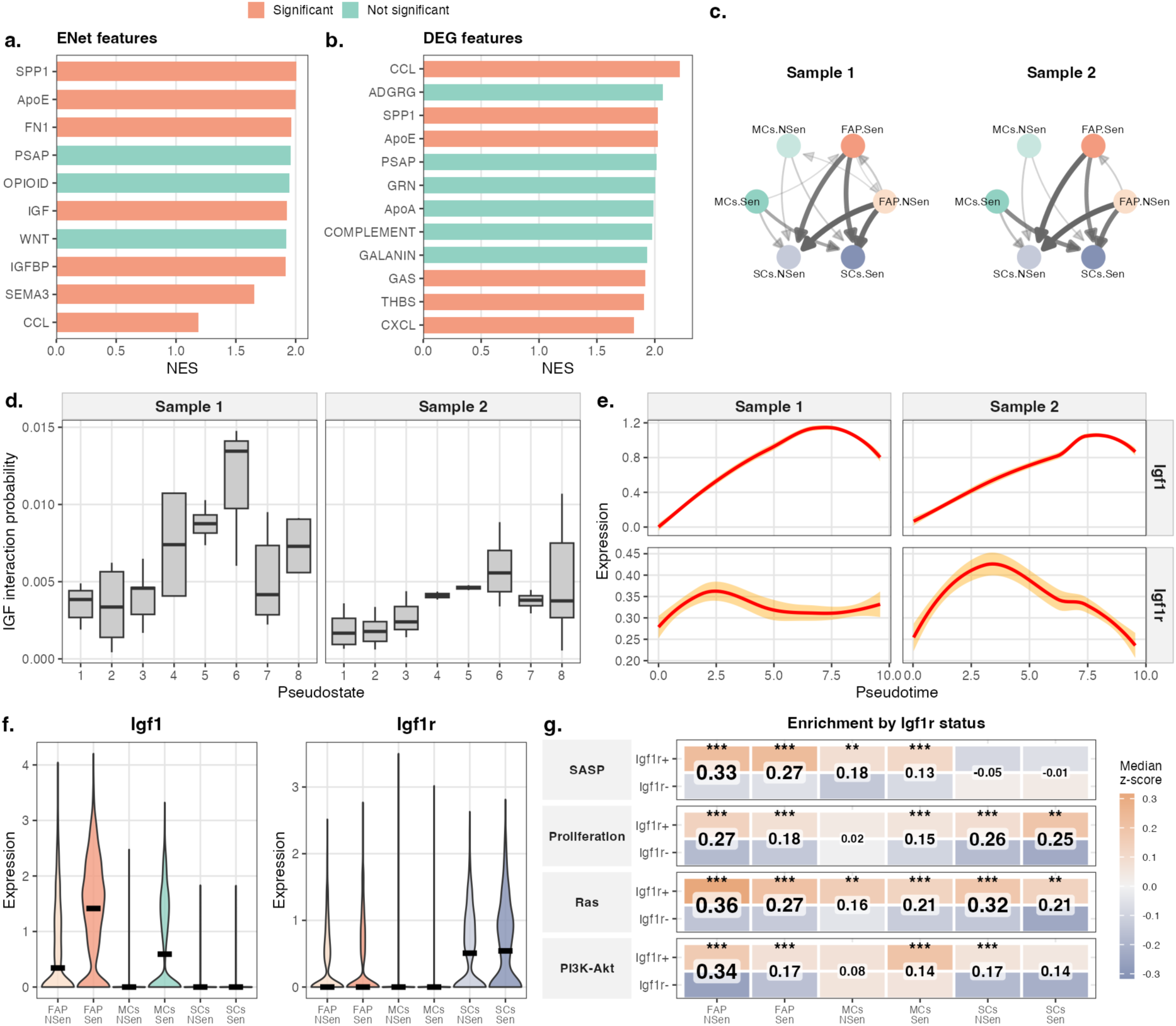
Senescence-associated cell–cell interactions and IGF signaling in senescence transduction. **(a)** Barplot showing normalized enrichment scores (NES) of interaction pathways from GSEA on ENet-prioritized features. Pathways are colored by significance of cell-cell interactions across cell types and senescence states (coral = significant interaction detected by CellChat, teal = not significant). **(b)** Same analysis performed on DEG-prioritized features (ranked by absolute log2 fold change). Several pathways (CCL, SPP1, ApoE) are identified by both methods, while others (FN1, IGF, IGFBP, SEMA3) emerge only through ENet modeling. **(c)** Circle plots illustrating IGF signaling-based cell-cell interaction networks in two biological replicates. Nodes represent cell type × senescence state combinations (FAPs, MCs, SCs crossed with Sen/NSen). Arrow direction indicates predicted sender to receiver based on ligand-receptor expressions; edge width corresponds to aggregated communication probability. **(d)** IGF-based cell-cell interaction probability across pseudostates (1–8). Interaction probability increases along pseudotime, peaks at late pseudostates (6–7), then declines in both biological replicates. **(e)** Expression trajectories of Igf1 (ligand) and Igf1r (receptor) along pseudotime in both biological replicates. Igf1 expression mirrors the interaction probability trajectory, peaking late; Igf1r peaks earlier and remains more stable. **(f)** Violin plots of Igf1 and Igf1r expression across cell types and senescence states. Igf1 is elevated in senescent FAPs and MCs (emitters); Igf1r is upregulated in SCs (receivers), consistent with directional IGF signaling from senescent cells to satellite cells. Black crossbars indicate medians. **(g)** Heatmap of pathway enrichment scores (SASP, Proliferation, Ras, PI3K-Akt) stratified by Igf1r expression status (Igf1r+ vs Igf1r-) across cell types and senescence states. Fill color indicates median z-score (normalized within each cell type × senescence combination); peach = higher enrichment, slate blue = lower. Cohen’s d effect sizes are displayed between paired rows, with text size proportional to effect magnitude. Significance: *p<0.05, **p<0.01, ***p<0.001 (Wilcoxon rank-sum test). Igf1r+ cells show elevated SASP and downstream signaling (Ras, PI3K-Akt), particularly in FAPs and senescent cells, while proliferation effects are more modest.

We next examined the functional correlates of IGF1 signaling in receiver cells, using *Igf1r* expression as a receptor proxy (**Fig. 4g**, **Extended Data Fig. 5**). In FAPs and MCs, *Igf1r*-positive cells exhibited significantly higher SASP enrichment scores (SenMayo). SCs showed no significant SASP enrichment shift with *Igf1r* status (**Fig. 4g**, **Extended Data Fig. 5a**). *Igf1r* expression was associated with increased proliferation scores, with significant effects in FAPs and SCs, but not in non-senescent MCs (**Extended Data Fig. 5b**). We then examined two canonical pathways downstream of IGF signaling^39^. Ras signaling pathway enrichment was significantly elevated in *Igf1r*-positive cells across all cell types and senescence states, with larger effect sizes in FAPs than in MCs (**Extended Data Fig. 5c**). PI3K-Akt signaling showed elevated enrichment in *Igf1r*-positive cells in FAPs, senescent MCs, and non-senescent SCs. Effects were not significant in non-senescent MCs or senescent SCs (**Extended Data Fig. 5d**). This indicates that in SCs, the main receivers of IGF signaling during secondary senescence, Ras signaling is constitutively activated in *Igf1r*-expressing cells regardless of senescence status. In contrast, PI3K–Akt signaling is conditionally activated in *Igf1r*-expressing non-senescent SCs.

Together, these results indicate that IGF signaling is directional and temporally staged, with *Igf1r* expression preceding *Igf1* along the senescence trajectory. The increase in inferred IGF communication when IGFBPs are computationally masked is consistent with IGFBP-mediated sequestration of IGF ligands. IGF reception correlates with senescence state, showing association with SASP gene expression in both non-senescent and senescent *Igf1r*-positive cells, particularly among FAPs and MCs. Finally, downstream pathway enrichment patterns vary by senescence state and cell type, suggesting that IGF signaling transduction engages context-dependent rather than uniform downstream effectors.

### Cross-context application reveals cell type-specific senescence accumulation and entropy-senescence coupling

To assess how the ENet classifier trained on single-cell data from young mice at 3 days post-injury (dpi) performs in other experimental contexts, we applied the model to bulk RNA-seq samples from the same study (GSE196613), which were generated from distinct biological samples. In this dataset, FACS-sorted cell populations representing different cell types and senescence states were sequenced using bulk RNAseq. The classifier achieved strong performance in fibro-adipogenic progenitors (FAPs, AUC = 0.94, n = 20) and myeloid cells (MCs, AUC = 0.90, n = 23), with AUC values ranging from 0.83 to 1.0 across conditions (Fig. 5a,b). Across all three cell types, the ENet model achieved robust classification of senescent samples in young mice at 3 days post-injury, the condition under which the model was originally trained using single-cell RNA-seq data. This observation underscores the senescence heterogeneity across age groups and post-injury time points. Although the trained model exhibits a degree of generalizability, its performance is highest under the specific conditions that match its training context, particularly for satellite cells (SCs). SCs showed more variable performance (overall AUC =0.83, n = 24), consistent with our held-out test results indicating that SCs represent the most challenging cell type for senescence classification. ENet percentile ranks showed clear separation between senescent and non-senescent cells across all the three cell types (Fig. 5c). Overall, despite the difference in sequencing strategy (single-cell RNA-seq versus bulk RNA-seq) and biological samples, these results corroborate ENet as an effective model for distinguishing SPiDER-based senescence.

**Figure 5.**
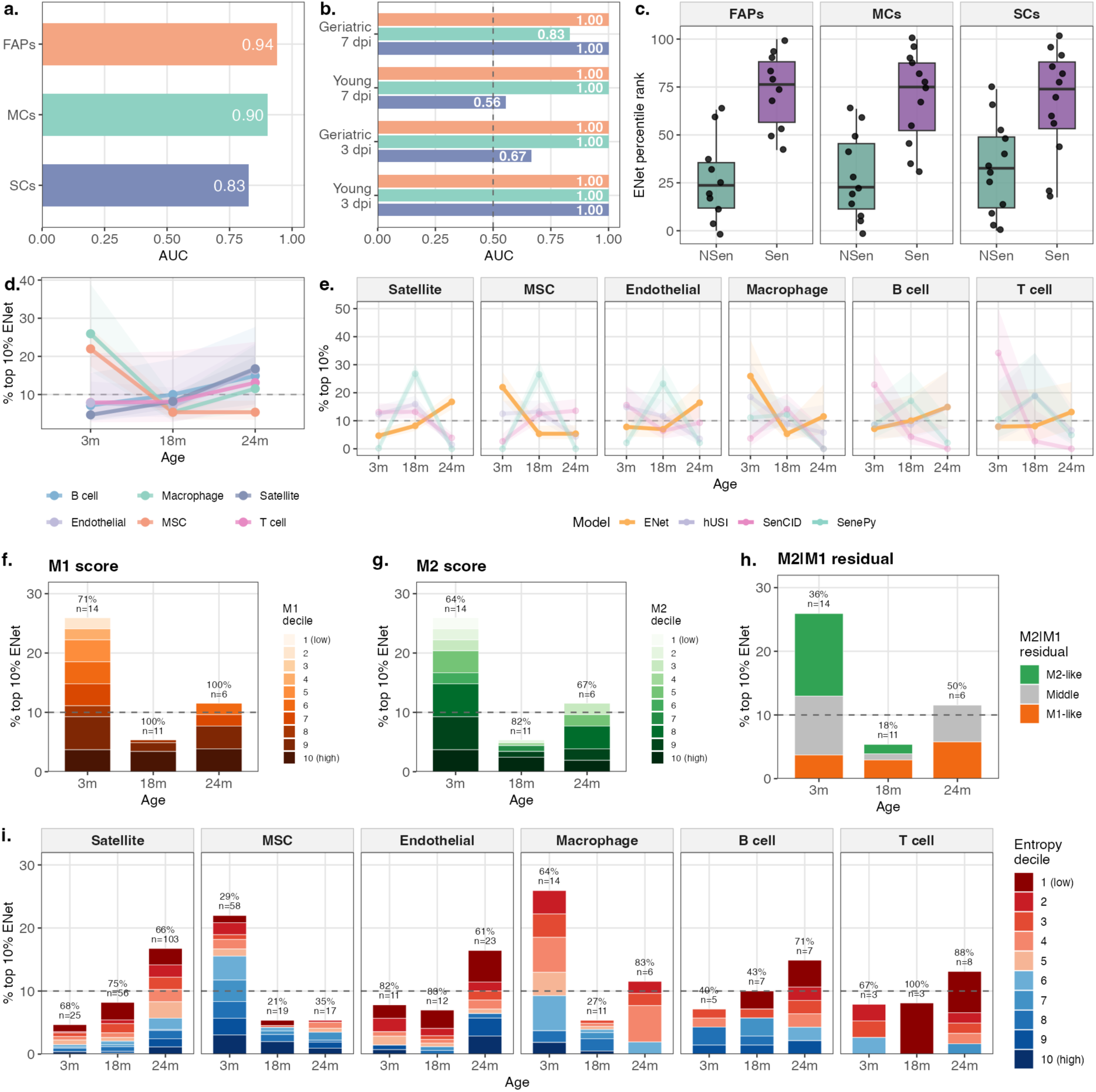
ENet senescence classifier performance across experimental contexts and cell type-specific aging patterns. **(a)** Classification performance (AUC) of the ENet senescence classifier applied to bulk RNA-seq samples from GSE196613, stratified by cell type. The classifier was trained on single-cell data from young mice at 3 days post-injury (dpi) and tested on bulk samples across conditions. AUC values are shown within bars. Colors indicate cell type (MCs, teal; FAPs, coral; SCs, slate blue). **(b)** AUC values stratified by age group (Young versus Geriatric) and injury timepoint (3 dpi versus 7 dpi) for each cell type. The dashed line indicates AUC = 0.5 (random performance). Colors are the same as panel a and indicate cell type. **(c)** Distribution of ENet percentile ranks in SPiDER-positive (Sen, purple) and SPiDER-negative (NSen, teal) cells from bulk RNA-seq samples, stratified by cell type. Percentile ranks were computed within each cell type. Box plots show median and interquartile range; individual samples shown as points. **(d)** Percentage of cells classified as senescent (ENet score above 90th percentile threshold) across ages (3, 18, and 24 months) in the Tabula Muris Senis limb muscle dataset. Shaded ribbons indicate 95% Wilson score confidence intervals. Dashed line indicates 10% reference (expected if random). Colors indicate cell type. **(e)** Comparison of four senescence prediction methods (ENet, hUSI, SenCID, SenePy) across cell types and ages. Each panel shows one cell type; y-axis indicates the percentage of cells exceeding the 90th percentile threshold for each method. **(f)** Distribution of senescent macrophages across M1 marker score deciles by age. Colors indicate decile. Text annotations show percentage of senescent macrophages in high M1 deciles (6 to 10) and total count. **(g)** Distribution of senescent macrophages across M2 marker score deciles by age. Format as in panel f. **(h)** Distribution of senescent macrophages by M1/M2 polarization balance. M2 scores were regressed against M1 scores, and residuals were computed for each macrophage. Positive residuals indicate M2-skewed cells; negative residuals indicate M1-skewed cells. Within each age group and residual sign, cells in the top 60% of absolute residual magnitude were classified as M2-like (green) or M1-like (orange); remaining cells were classified as intermediate (gray). Text annotations show percentage in intermediate category and total count. **(i)** Distribution of senescent cells across transcriptional entropy deciles (1 = lowest, 10 = highest entropy) for each cell type and age. Stacked bars show percentage of total cells classified as senescent within each entropy decile. Text annotations indicate the percentage of senescent cells in low entropy deciles (1 to 5) and total senescent cell count. Dashed line indicates 10% reference.

We applied the ENet classifier to single-cell RNA-seq data from the Tabula Muris Senis (TMS) project, analyzing limb muscle samples from mice at 3, 18, and 24 months across six cell types (satellite cells, n = 1,833; mesenchymal stem cells, n = 935; endothelial cells, n = 453; macrophages, n = 311; B cells, n = 187; T cells, n = 136). Cells exceeding the 90th percentile ENet score threshold were classified as senescent (Fig. 5d). The percentage of senescent cells increased with age in satellite cells (4.67% to 16.7%), endothelial cells (7.80% to 16.4%), B cells (7.14% to 14.9%), and T cells (7.89% to 13.1%). Mesenchymal stem cells (MSCs) and macrophages showed the opposite pattern, decreasing from 22.0% to 5.36% in MSCs and from 25.9% to 11.5% in macrophages. Comparison with three alternative methods (hUSI, SenCID, SenePy) revealed divergent age-dependent trends (Fig. 5e). While ENet detected age-associated increases in four cell types, alternative methods showed stable or decreasing trends. For macrophages, ENet and hUSI detected decreased senescence with age, while SenCID and SenePy showed increases from 3 to 18 months, followed by zero detection at 24 months.

Given the decline in predicted senescent macrophages with age, we examined their phenotypic characteristics using M1 (pro-inflammatory) and M2 (anti-inflammatory) marker signatures^40^. Senescent macrophages showed elevated expression of both M1 (0.266 versus 0.115, P = 3.5 × 10^-12) and M2 markers (0.131 versus 0.064, P = 3.4 × 10^-7), with 71% to 100% in high M1 deciles and 64% to 82% in high M2 deciles across ages (Fig. 5f,g). Using M2 versus M1 residual-based classification (Fig. 5h), the polarization balance shifted with age, from 50% M2-like and 14% M1-like at 3 months (n = 14) to 27% M2-like and 55% M1-like at 18 months (n = 11), and finally 0% M2-like and 50% M1-like at 24 months (n = 6). All senescence prediction methods detected elevated M1 and M2 markers in senescent macrophages at young ages (Extended Data Fig. 6a-c), suggesting senescence signatures overlap with macrophage activation states. The age-dependent decline in detected senescence for ENet and hUSI may partly reflect the loss of polarized macrophages in aged muscle.

To test whether the reduced transcriptional entropy in senescent cells observed in the original dataset holds across cell types and aging contexts, we computed Shannon entropy for each cell in the TMS dataset (Fig. 5i). Three cell types showed significantly lower entropy in senescent cells, namely satellite cells (9.50 versus 10.0, P = 1.09 × 10^-21), endothelial cells (9.41 versus 9.78, P = 1.90 × 10^-4), and T cells (9.38 versus 10.1, P = 2.64 × 10^-4). In contrast, MSCs showed significantly higher entropy in senescent cells (10.8 versus 10.4, P = 2.73 × 10^-12), while macrophages (P = 0.084) and B cells (P = 0.186) showed no significant difference. In satellite cells, 66% to 75% of senescent cells fell within low entropy deciles (1 to 5) across all ages, whereas MSCs showed the opposite pattern, with only 21% to 35% in low entropy deciles. The high entropy in MSC senescent cells, combined with their enrichment at young ages, suggests these cells may differ functionally from the low-entropy senescent cells observed in other cell types. Whether this more plastic senescence state reflects transient versus persistent senescence states, or cell-type-specific senescence programs, remains to be investigated. The alternative senescence prediction methods did not show the same low-entropy enrichment (Extended Data Fig. 6d). In satellite cells, only 18% to 26% of senescent cells identified by hUSI, SenCID, and SenePy fell within low entropy deciles, compared with 66% to 75% for ENet. ENet scores were negatively correlated with entropy, while alternative methods showed no consistent relationship (Extended Data Fig. 6e), indicating that ENet preferentially identifies transcriptionally constrained cells.

## Discussion

A central challenge in senescence biology is whether cellular senescence states can be reliably inferred from transcriptomic data when grounded in experimentally validated labels, and whether data-driven models provide advantages over marker-centric definitions. Here, using SPiDER-β-gal–labeled senescent cells as in vivo ground truth, we demonstrate that multigene, jointly weighted models substantially outperform marker-centric and enrichment baselines, while revealing structured, cell-type–specific senescence programs and candidate signaling axes involved in secondary senescence.

We first observed pronounced discordance among classical marker sets by benchmarking against the experimentally determined β-gal activity. SenMayo enrichment scores increased in SPiDER⁺ cells, whereas GenAge and CellAge scores decreased, despite all being widely used to annotate senescence. This divergence likely reflects the distinct biological emphases of these gene sets: SenMayo is enriched for inflammatory and SASP-related genes, whereas GenAge and CellAge aggregate broader, heterogeneous associations with aging and senescence. Importantly, apparent concordance between these scores can be recovered when SenMayo enrichment itself is used to define senescence status, illustrating a form of circular evaluation. A composite ratio (CellAge/SenMayo) partially reconciled these opposing signals and improved discrimination relative to individual gene sets. However, improved performance likely results from the anti-correlation of CellAge and SenMayo rather than providing additional biological insight.

Motivated by these limitations, we applied regularized machine-learning models that jointly weight transcriptome-wide features. Across sample-held-out evaluations, all models outperformed marker- and enrichment-based baselines, with the largest relative gains observed in satellite cells, the most challenging lineage for all approaches. This improvement is consistent with the ability of multivariate models to integrate weak but coherent signals that are lost in additive scoring schemes. ENet-prioritized features showed only partial overlap with conventional DEGs and instead formed correlated modules spanning stress response, extracellular matrix remodeling, metabolic rewiring, and cell-cycle checkpoint regulation, reinforcing the view of senescence as an emergent, systems-level state rather than the dysregulation of a small set of canonical genes.

A potential caveat in transcriptome-based classification is “cell-type leakage”, whereby lineage identity rather than cellular state drives predictive performance. Canonical senescence readouts such as Cdkn2a, Cdkn1a, and SA-β-gal activity exhibit strong baseline differences across cell types, complicating cross-lineage comparisons^8^. However, stratified evaluation within cell types, combined with balanced Sen-NSen sampling during training, argues against a purely identity-driven explanation in our study. Nonetheless, future extensions incorporating explicit deconfounding strategies—such as residualization of lineage signatures, adversarial learning objectives, or cross-lineage transfer tests—may further improve generalizability.

Trajectory analyses derived solely from ENet-prioritized features recovered late branches enriched for SPiDER⁺ cells without using senescence labels as supervision, indicating that the ENet model representation captures progression information intrinsic to the transcriptome. These trajectories exhibited lineage-specific organization, representing pathway-dependent heterogeneity in senescence programs^26^. Notably, in satellite cells, advanced non-senescent branches were enriched for cell–cell communication and signal-transduction pathways, suggesting a role for paracrine signaling in driving secondary senescence. Integrating trajectory inference with ML-based prediction yielded a pseudotime-like score that may be particularly sensitive for detecting progressive or early senescent states in vivo—an important advantage given the rarity and transient nature of fully senescent cells during natural aging.

Entropy analyses provided an orthogonal validation of the classification framework. Across FAPs, MCs, and SCs in the injury model, senescent cells exhibited reduced transcriptional entropy relative to non-senescent cells, a pattern that was recapitulated in bulk RNA-seq data. Non-senescent cells from injured tissue displayed elevated entropy compared to uninjured controls, consistent with regeneration-associated transcriptional diversification. Because senescent and non-senescent cells were compared within the same injury context, these analyses isolate senescence-associated effects from injury-induced transcriptional changes. Together, these results suggest that transcriptomic entropy may serve as an independent metric for evaluating senescence classification.

Cell–cell communication analyses converged on the IGF signaling axis as a candidate mediator of secondary senescence. Ligand expression was predominantly observed in senescent FAPs and macrophages, whereas receptors were enriched in satellite cells, with Igf1 expression increasing along pseudotime in emitters and Igf1r elevated in receivers. Computationally masking IGF-binding proteins (IGFBPs) increased inferred interaction probabilities, consistent with their known role in modulating IGF bioavailability and signaling range^3,37^. These findings align with prior studies demonstrating that IGF-I promotes senescence in multiple cell types by increasing reactive oxygen species and activating the p53–p21 pathway^41^. The association between Igf1r expression and elevated SASP, RAS, and PI3K–Akt pathway activity supports a functional link between IGF reception and senescence phenotypes^42^. Mechanistically, IGF-driven PI3K–AKT signaling regulates FOXO3 phosphorylation and nuclear exclusion, impairing transcriptional activation of protective programs and promoting senescence. Recent evidence showing systemic rejuvenation upon FOXO3 dephosphorylation reinforces the plausibility of this cascade as a regulator of senescence in mice and monkeys^43^. Nevertheless, given the injury context of the dataset, IGF signaling may also reflect regenerative processes, and targeted perturbation studies will be required to disentangle causality.

Application of the ENet model to the Tabula Muris Senis aging dataset revealed age-associated accumulation of senescent states in satellite cells, endothelial cells, and lymphocytes, consistent with established roles of senescence in tissue aging^23,44,45^. In contrast, mesenchymal stromal cells and macrophages showed decreased predicted senescence with age. For macrophages, this trend coincided with the loss of M2-polarized cells, and senescent classifications were associated with elevated expression of both M1 and M2 markers across methods, suggesting that transcriptomic senescence signatures may partially capture activated immune states. In mesenchymal stromal cells, entropy analyses revealed an opposite pattern, with ENet-identified senescent cells exhibiting elevated entropy and being enriched at younger ages. This raises the possibility that the model detects a transient, developmentally associated senescence-like state in this lineage, though alternative explanations, including cell-type–specific programs or classifier limitations, cannot be excluded.

Several limitations should be noted. The primary dataset captures a single acute injury time point, and senescence programs and paracrine interactions may differ across tissues, stressors, or chronic aging contexts. SPiDER-β-gal provides a live-cell–compatible readout of β-galactosidase activity but is neither fully sensitive nor specific for senescence, and our classifiers therefore predict SPiDER positivity rather than an absolute senescence state. While sample-held-out evaluation and cross-platform validation on bulk RNA-seq from different biological samples strengthen internal validity, additional experimentally labeled datasets will be required to assess cross-context portability. Finally, percentile-based thresholding assumes a fixed proportion of senescent cells within each context, facilitating within-tissue comparisons but precluding direct estimation of absolute senescence prevalence across tissues or datasets.

In summary, experimentally grounded single-cell analyses support the use of multigene, jointly weighted models over marker-centric approaches for senescence classification. Our framework recovers cell-type–specific progression trajectories, identifies IGF signaling as a plausible axis of secondary senescence in regenerating muscle, and provides a scalable route from classification to mechanistic hypothesis generation. Extension of this approach to human tissues, longitudinal in vivo settings, and targeted perturbation of candidate pathways will be important next steps toward translating senescence-informed models into therapeutic insight.

## Methods

### Biological material, labeling, and library construction

The main dataset (GSE196613) used here was obtained from the senescence atlas research generated by Moiseeva et al. (2023)^15^, targeting approximately 5,000 cells per run. In brief, each dataset was produced from two biological replicates of young mice (3-6 months old) subjected to senescence induction. Intramuscular injections of CTX (Latoxan, L8102; 10 µM) were administered to induce senescence in skeletal muscle. Mice were euthanized at three days post-injury (d.p.i.), after which muscles were dissected, flash-frozen in liquid-nitrogen-cooled isopentane, and stored at −80 °C until analysis. Cell sorting was conducted on CD45-positive and CD45-negative populations using PE-Cy7-conjugated anti-CD45 antibodies (Bio-Legend, 103114) and senescent cells were separated using SPiDER-β-gal (SPiDER) markers. Following cell sorting via the BD FACS Aria II system, scRNA-seq libraries were prepared with the Chromium Single Cell 3′ GEM, Library & Gel Bead Kit v3 (10x Genomics, PN-1000075). A total of 21,449 cells were collected, comprising three primary cell types: myeloid cells (MCs), fibroadipogenic progenitors (FAPs), and satellite cells (SCs). The corresponding bulk RNA-seq data, provided as the trimmed mean of M-values (TMM)-normalized counts per million (CPM) value matrices from dedicated cell types and senescent biological samples after sorting, were directly obtained from the same study and used in this work to validate transcriptomic entropy observed in single-cell RNA-seq data. The high sequencing depth of bulk RNA-seq supports robust entropy estimation. Detailed experimental procedures and preprocessing steps are available in the original publication^15^. Gene-level TPM values (Transcripts Per Million) from the same bulk RNA-seq data, generated using the same nf-core bulkRNAseq pipeline and procedures^46^ as in the original publication^15^, were used as input to predict senescence score using our classifier.

The in vitro dataset (GSE226225)^22^ is applied to validate the enrichment relationship between SenMayo and CellAge. In brief, WI-38 human diploid fibroblasts were cultured under standard conditions and subjected to three senescence-inducing protocols: replicative exhaustion (PDL > 50), ionizing radiation (10 Gy), and etoposide treatment (50 μM for six days followed by recovery). Etoposide (ETO) is an anti-cancer chemotherapy drug. Time-course samples were collected post-ETO treatment, and cell cycle synchronization was achieved using low-serum media. Senescence was validated by SA-β-Gal staining and BrdU incorporation assays, using established protocols and imaging or plate reader analysis. Single-cell RNA libraries were prepared using 10x Genomics Chromium kits, with two replicates per senescence model and one per ETO time point. Approximately 7,000 cells per sample were processed to generate gel bead-in-emulsions (GEMs), enabling RNA barcoding and reverse transcription. After cDNA synthesis and quality assessment, libraries were sequenced on an Illumina NovaSeq 6000, yielding 45,000–100,000 reads per cell.

Single-cell RNA-seq data from the Tabula Muris Senis (TMS) project to explore the senescence classification between our ENet model and others’ models, were obtained from the work of Lagger et al.^47^. Limb muscle samples from mice at 3, 18, and 24 months were analyzed across six cell types (satellite cells, mesenchymal stem cells, endothelial cells, macrophages, B cells, T cells). Cell type and age groups with fewer than 20 cells were excluded.

### Preprocessing, quality control, and normalization

We start from the processed data in the two datasets^15,22^. For the in vivo dataset, reads were aligned to the mouse genome (mm10, GENCODE vM23) using STARsolo. Downstream analysis was conducted in R using Seurat and DoubletFinder, with standard quality filtering and integration. Low-quality cells and doublets were excluded. We log-normalized the dataset using the Seurat package^48^ (v4.3.0.1). In brief, feature counts per cell were normalized by dividing by the total counts and multiplying by a scaling factor of 1e6, followed by a natural log transformation using log1p. Differentially expressed genes were identified using the FindMarkers function with the Wilcoxon Rank Sum test. A minimum expression threshold of either 25% (senescence DEGs) or 10% (trajectory lineage DEGs) of cells in either group was applied, and both upregulated and downregulated genes were included. Unless otherwise specified, a default log fold change threshold of 0.25 was used. Initially, there were 21,449 cells in total; we only kept the 3 major cell types of 21,239 cells in total with FAPs of 9,841 cells, MCs of 8,693 cells and SCs of 2,705 cells. For the in vitro dataset, 10x sequencing libraries were processed using the Cell Ranger pipeline (v5.0.0) to generate feature-barcode matrices, with reads mapped to the GRCh38 reference genome.

Quality-controlled single-cell data were analyzed using Seurat (v4.1.0), filtering cells based on mitochondrial RNA content, transcript counts, and gene expression thresholds. One low-quality control sample was excluded. Samples were normalized, highly variable genes selected, and datasets integrated using Seurat’s recommended pipeline. In the end, we got 5,585 cells in the control condition; 2,300 cells in the replicative senescence (RS) condition, 2,184 cells in the ionizing radiation (IR) condition; 5,834 cells in the etoposide (ETO) day 0; 3,033 cells in the etoposide (ETO) day 10.

### Single-marker baselines, curated set enrichment, and AUC calculation

For single-marker baseline analysis, we used p16 (*Cdkn2a*), p21 (*Cdkn1a*), and *Tgfb1*^17^. p16 and p21 are widely recognized senescence markers, commonly assessed alongside SA-β-Gal activity^8^. Although *Tgfb1* is not a direct marker of senescence, it plays a pivotal role in inducing the senescent phenotype through multiple pathways, including activation of p53/p21 and upregulation of p16 in specific contexts. Elevated *TGFB1* expression or activity is therefore associated with senescence across various cell types and disease states, making it a relevant factor in this analysis^49^. Normalized expression levels of these markers in SPiDER-labeled senescent groups are presented.

In addition to individual markers, senescence marker sets were curated from established sources, and enrichment analysis was conducted based on the marker sets. Raw counts are recommended to be used as input for enrichment score calculation. A modified “enrichIt” function from the escape package (v1.4.0) was used, setting ‘groups=1000’ and ‘ssgsea.norm=FALSÈ to maintain reproducibility. The function “enrichIt” implements a single-cell adaptation of single-sample gene set enrichment analysis (ssGSEA^50^) based on gene set variation analysis (GSVA), providing gene-set scores on a per-sample basis in an unsupervised manner. This approach enabled detailed scoring for individual cells in line with senescence marker enrichment.

To evaluate the discriminatory power of individual gene expression or gene set enrichment scores in distinguishing SPiDER-labeled non-senescent (NSen) and senescent (Sen) cells, we performed receiver operating characteristic (ROC) analysis using the pROC package (v1.18.4) in R (v4.1.3). For each gene or enrichment score, we extracted the corresponding numeric values across single cells and assigned binary labels (0 for NSen, 1 for Sen) based on SPiDER classification. ROC curves were generated using the roc() function, and the area under the curve (AUC) was computed using the auc() function.

### Differential expression (DE) and ML feature importance baseline

Differentially expressed genes (DEGs) were identified using the Seurat package (v4.3.0.1)^51^, as described in the preprocessing section. As a baseline, DEGs with Bonferroni-adjusted p-values below 0.05 were selected. To further refine the set based on effect size, we applied an elbow point detection method to the absolute values of average log₂ fold changes (avg_log2FC), identifying a threshold that captures genes with substantial expression changes. A similar elbow test was applied to determine the cutoff for important features in the Elastic Net (ENet) model, based on the ranked coefficients. Specifically, we used the uik() function from the inflection package (v1.3.6) in R to compute the knee point—where the rate of change in feature importance sharply declines, indicating diminishing returns.

The resulting DEG set from the NSen versus Sen comparison in the training sample was used as a data-driven positive baseline for evaluating the classification performance of alternative gene sets and methods in distinguishing SPiDER-labelled senescent states.

To evaluate potential cell-type confounding, we performed an enrichment analysis of ENet-important features against cell-type–specific marker genes (**Extended Data Fig. 2e,f**). First, we identified cell-type markers by combining both samples after applying the same criteria used for defining significant DEGs. Next, we removed all senescence-associated genes from these marker sets—both in a cell-type–specific manner and in a cell-type–agnostic manner—to obtain senescence-unrelated marker gene sets. These filtered marker sets were then used to test enrichment among the ENet-important features.

### Machine-learning classifiers and feature selection

No batch effect was observed across the two biological replicates, as indicated in the original publication^15^, allowing them to remain independent for machine learning purposes. Sample #2 (n=11,612 total cells, 7,749 senescent, and 3,863 non-senescent) was used for training, while sample #1 (n=9,627 total cells, 6,157 senescent, and 3,470 non-senescent) was used for validation due to its smaller cell count. Gene expression counts are normalized per cell and scaled across cells within training and test datasets separately, with the same scaling size factor based on the training dataset. When applying the trained models to new datasets, features are scaled using the same scaling factors derived from the training dataset, unless otherwise specified. The processed gene expression counts serve as feature values. The top 2000 variable genes in the training sample were selected based on a 1:5 feature-to-sample ratio. Variable features were identified through the “vst” method, where a local polynomial regression (loess) line was fitted to the log-variance vs. log-mean relationship, after which feature values were standardized based on the observed mean and expected variance. To classify senescent versus non-senescent cells based on transcriptome profiles, three models were evaluated: Elastic Net (glmnet), Support Vector Machine with a linear kernel (svmLinear), and Random Forest (rf) via the caret package (v6.0-94). Cross-validation (10-fold) on the training replicate was used for hyperparameter tuning and feature selection. Model performance was assessed using sample #1, independent of cell types. To identify optimal feature subsets for classification for the Elastic Net model, we performed Recursive Feature Elimination (RFE) using the caret package (v6.0-94)^52^ in R. For Elastic Net, a grid search was applied over a range of alpha (0 to 1) and lambda (0.001 to 0.1) values. RFE was conducted using 10-fold cross-validation (method = “cv”, number = 10) with parallel processing enabled. Feature subset sizes ranged from 5 to 2000. The rfeControl() function was used to define the resampling strategy and control parameters, and model performance was assessed on the final resampling results.

### Comparison with other machine-learning classifiers

For method comparison, the ENet classifier was applied to the log-transformed library-size-normalized gene counts of bulk RNA-seq samples from GSE196613 across three cell types (FAPs, MCs, SCs), two age groups (young, geriatric), and two timepoints after injury (3 dpi, 7 dpi). Classification performance was evaluated using AUC with SPiDER SA-β-gal labeling as ground truth. ENet percentile ranks were computed within each cell type.

The ENet classifier trained on GSE196613 was applied to TMS data without retraining. Cells exceeding the 90th percentile ENet score within each cell type were classified as senescent. Alternative methods (hUSI, SenCID, SenePy) were computed using published implementations^13,14,20^ with the same thresholding approach. Feature names originating from human datasets were converted to their corresponding mouse orthologs by refering the conversation file from the Mouse Genome Informatics (MGI) database^53^.

### Trajectory inference

Pseudotime trajectory analysis was conducted using Monocle 2 (v2.22.0)^54^ by using the important features from the ENet model defined by the feature coefficient more than the threshold based on the elbow test. Monocle 2 orders single cells along a trajectory to model dynamic biological processes, such as cell differentiation. The analysis involves dimensionality reduction using Uniform Manifold Approximation and Projection (UMAP) and trajectory inference. Pseudotime is defined as the relative distance between each cell and the root cell along the inferred trajectory. The root cells of the trajectory were identified as the cell branch that achieved the highest entropy.

### Transcriptional entropy

To quantify transcriptomic entropy in bulk RNA-seq data or the TMS data, we computed Shannon entropy across all expressed genes for each sample. Zero values were excluded to avoid undefined logarithmic operations. Entropy was then calculated using the standard Shannon formula: *H* =− ∑ *p_i_ log*2(*p_i_*), where (*p_i_*) is the normalized expression proportion of gene ( i ) in a given sample.

To assess transcriptional heterogeneity at the single-cell level, we calculated Shannon entropy across the full transcriptome using normalized expression data. The Seurat object was converted to a SingleCellExperiment object, and entropy was computed using the perCellEntropy() function from the TSCAN package (v1.32.0)^55^ in R (v4.1.3), applied to all expressed genes. This approach captures global transcriptional diversity per cell, providing a quantitative measure of cellular plasticity and complexity.

### Gene ontology enrichment analysis

Gene Ontology (GO) enrichment analysis was performed to characterize biological processes associated with transcriptional changes for either ENet important features or along pseudotime-defined branches for each cell type. Differential expression statistics were generated for trajectory state comparisons in MCs, SCs, and FAPs. For each comparison, genes were ranked in descending order by feature importance metrics or fold change and used as input for gene set enrichment analysis (GSEA).

GSEA was carried out using the ‘gseGO()’ function from clusterProfiler (v4.2.2)^56^ with the following parameters: ‘keyType = “SYMBOL“’, ‘OrgDb = org.Mm.eg.db’, ‘ontology = “ALL“’, ‘minGSSize = 20’, ‘maxGSSize = 1000’, and ‘pvalueCutoff = 1’ to retain all pathways prior to downstream filtering. Enrichment analysis was performed separately for each trajectory comparison across MCs, SCs, and FAPs. To reduce redundancy among enriched GO terms, results were simplified in the website of REVIGO^57^, Simplified enrichment results were used for visualization and downstream interpretation.

### Ligand–receptor inference with CellChat

Cell–cell communication networks were inferred using the CellChat R package^58^. Preprocessed single-cell RNA-seq data were used as input, with cells grouped by annotated cell types. The CellChat mouse database^59^ (CellChatDB.mouse) was applied as the reference ligand–receptor database. Overexpressed genes and interactions were identified, followed by the computation of communication probabilities (triMean method) and filtering based on a minimum of 10 cells per group. Communication probabilities were aggregated at the signaling pathway level, and both the number and strength of interactions were quantified and visualized using circle plots. Network centrality analysis was performed to identify dominant sender and receiver cell populations for each signaling pathway. The pseudostated interaction probability was calculated by grouping cells into the same pseudostate, defined by pseudotime bins, and then running cell–cell interactions based on cell type and senescence state within each pseudostate. We performed pathway enrichment analysis using features ranked by model-specific importance metrics: feature coefficients for Elastic Net (ENet) and Support Vector Machine (SVM) models, and mean decrease in Gini index for Random Forest (RF). Ranked features were tested against 193 signaling pathways from CellChatDB using the fgsea package (v1.20.0). This analysis identified enriched interaction pathways for each machine learning model, from which significant cell type– and senescence state–specific interaction pathways were also reported.

### Pathway scoring and proliferation indices

Mouse gene sets representing IGF downstream signaling and muscle cell proliferation were obtained from MSigDB^60,61^. For each single cell, pathway activity and proliferation indices were quantified using single-sample Gene Set Enrichment Analysis (ssGSEA), implemented via the enrichIt() function from the escape package (v1.4.0). The resulting enrichment scores reflect the relative activation of each gene set across individual cells.

### Macrophage polarization scoring

M1 and M2 polarization scores were computed by the Seurat function of “AddModuleScore()” using gene signatures from Oshi et al.^40^ as average expression of signature genes. To classify relative M1/M2 balance, M2 scores were regressed against M1 scores across all macrophages. Cells with positive residuals in the top 60% of absolute magnitude were classified as M2-like, negative residuals in the top 60% as M1-like, and the remainder as intermediate.

### Statistics

All statistical comparisons between two groups were performed using two-tailed t-tests, assuming equal variance unless otherwise specified. For comparisons involving more than two groups, analysis of variance (ANOVA) was applied to assess overall significance, followed by appropriate post hoc tests where relevant. Statistical analyses were conducted in R (v4.1.3), and significance thresholds were set at p < 0.05 unless otherwise noted. Confidence intervals (95%) for the proportion of senescent cells were computed using the

Wilson score formula: 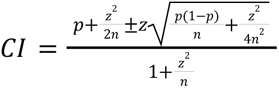, where p is the observed proportion, n is the sample size, and z = 1.96 for 95% confidence. The Wilson method was preferred over standard Wald intervals 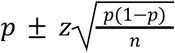 because Wald intervals can produce values outside [0,1] and have poor coverage when proportions are near 0 or 1 or when sample sizes are small.

## Data and code availability

The source data and plotting objects supporting all main and extended data figures are available from the BioStudies database under accession number S-BSST2347.

The primary single-cell RNA-sequencing data used for model training and evaluation are available from the Gene Expression Omnibus under accession GSE196613. Independent WI38 cell line datasets are available under GSE226225. The Tabula Muris Senis dataset used for model application is available at https://doi.org/10.6084/m9.figshare.19590256.All analysis scripts and processed metadata generated in this study are available at https://github.com/donertas-group/senClassification_SPiDER.git, enabling full reproducibility of the results reported in this manuscript.

## Acknowledgement

The project is funded by the Carl-Zeiss-Stiftung (P2021-00-007, HMD). We acknowledge the core facility of Life Science Computing in the Leibniz Institute on Aging – Fritz Lipmann Institute (FLI) for their management of high-performance computing resources that supported this work.

## Author contributions

M.D. conceived the study and supervised the project. M.D. and J.L. designed the computational framework. J.L. implemented the bioinformatics pipelines, performed data curation, and statistical analyses with conceptual contributions from all authors. J.L. and M.D. created visualizations and wrote the original draft. All authors contributed to reviewing and editing the manuscript and approved the final version.

## Supplementary Materia

**Extended Data Figure 1:**
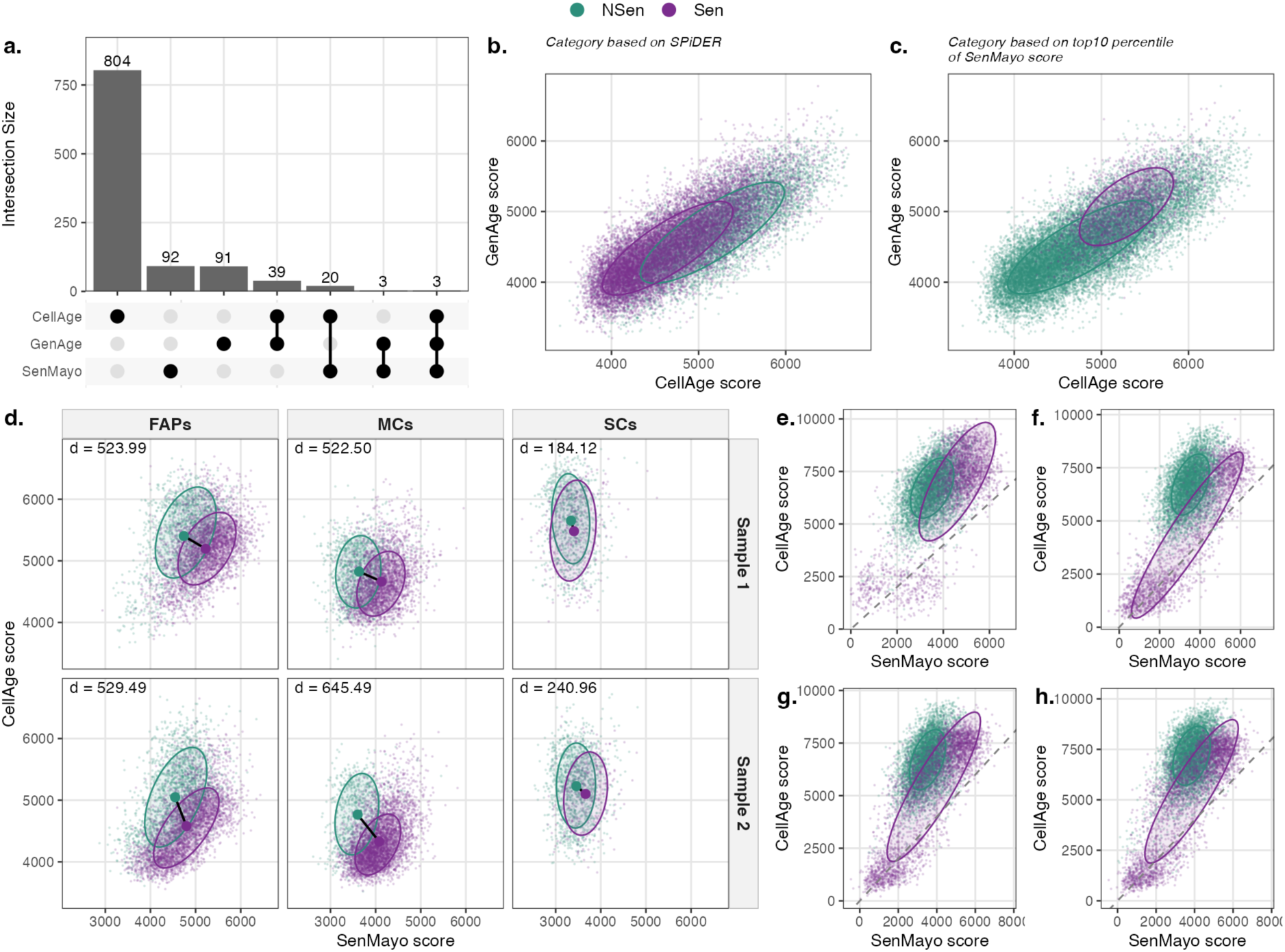
Curated senescence marker sets outperform single markers. **a)** UpSet plot showing the overlap between the SenMayo, GenAge, and CellAge gene sets. CellAge contained the largest number of unique genes. **b–c)** Scatter plots showing CellAge and GenAge enrichment scores. Cells colored by **(b)** SPiDER classification or **(c)** top decile of SenMayo enrichment. Ellipses indicate 68% data coverage for each group. Purple: senescent (Sen); teal: non-senescent (NSen). **d)** Scatter plots of SenMayo versus CellAge enrichment scores stratified by cell type (FAPs, MCs, SCs) and sample replicate. SPiDER-positive cells showed consistent enrichment patterns across cell types. Text annotation d indicates the euclidean distance between centroids. **e–h)** Validation in an independent WI-38 dataset (GSE226225). Enrichment of SenMayo versus CellAge in experimentally induced senescence models. Human diploid WI-38 fibroblasts were cultured under standard conditions and subjected to **(e)** replicative senescence (RS; population doubling level > 50), **(f)** ionizing radiation-induced senescence (IR; 10 Gy, harvested 10 days post-irradiation), or **(g,h)** etoposide-induced senescence (ETO; 50 μM for six days followed by four days recovery). Panels **(e–g)** compare senescent cells to proliferating controls (PDL 24), while **(h)** shows ETO day 10 vs day 0. Dashed diagonal lines indicate y = x. Senescence induction was confirmed by senescence-associated β-galactosidase (SA-β-Gal) staining and reduced BrdU incorporation. Proliferating controls/ETO day0 cells are shown in teal, senescent conditions in purple.

**Extended Data Figure 2:**
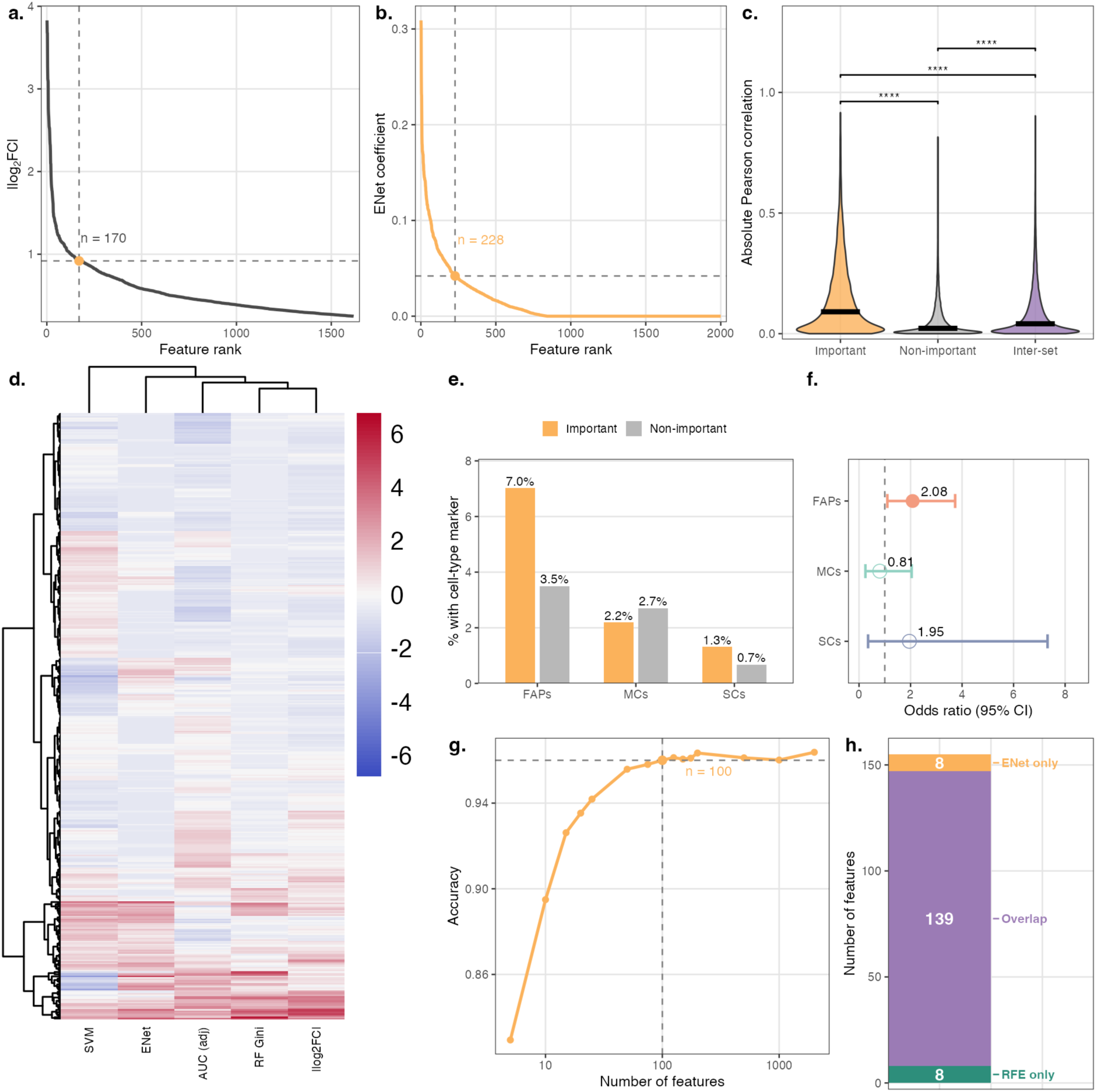
Feature selection and ML model characterization. **(a)** Feature ranking by absolute log₂ fold-change identifies 170 DEGs at the elbow threshold (dashed lines). (**b)** ENet model coefficient ranking identifies 228 important features at the elbow threshold. **(c)** Pairwise correlation distributions among important features (n=228), non-important features, and inter-set comparisons. Important features show significantly stronger internal correlations compared to non-important (p < 2.2×10⁻¹⁶) or inter-set correlations, suggesting coordinated expression patterns. **(d)** Heatmap of feature-importance metrics for the 416 features shared across all methods. A total of 2,000 variable features were identified by machine-learning models (ENet, RF, SVM), and 1,440 DEGs were selected based on ‘logfc.threshold = 0.25’ and ‘min.pct = 0.25’. Feature-level AUC values were computed for all 26,643 genes, and intersecting these sets yielded the 416 common features visualized here. Displayed metrics include ENet and SVM coefficients, mean decrease in Gini index from the RF model, and direction-adjusted AUC or absolute log₂ fold change for DEGs. Features are hierarchically clustered based on similarity across all metrics. **(e)** Senescence unrelated cell-type marker enrichment among ENet-important features. Grouped bar chart shows the percentage of features with cell-type markers for important (golden) vs non-important (gray) feature sets across FAPs, MCs, and SCs. **(f)** Forest plot of odds ratios (95% CI) for cell-type marker enrichment. Filled circles indicate significant enrichment (p < 0.05). FAPs show significant enrichment of cell-type markers among important features (OR = 2.08, p = 0.017), while MCs (OR = 0.95, p = 0.83) and SCs (OR = 1.95, p = 0.24) do not reach significance. **(g)** Recursive feature elimination (RFE) identifies optimal model size at 100 features (dashed line), achieving 96% accuracy while reducing feature count by 56% from the full 228-feature set. **(h)** Feature selection convergence: 147 features identified by RFE show substantial overlap (139 features, 95%) with top ENet-ranked features, demonstrating reproducibility across resampling schemes. Colors indicate RFE-only (teal, n=8), overlap (purple, n=139), and ENet-only (golden, n=8) features.

**Extended Data Figure 3:**
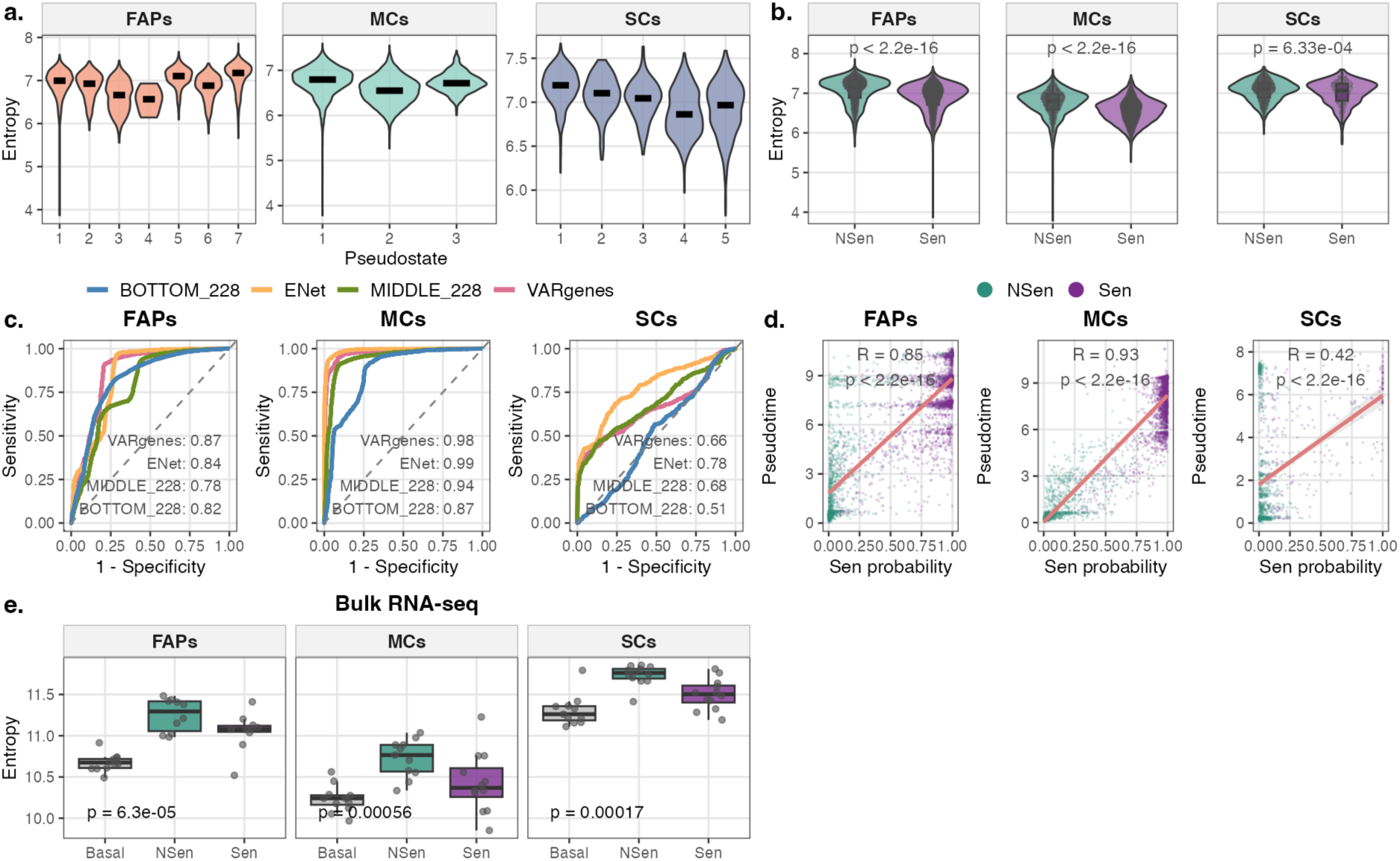
ML-prioritized features reconstruct senescence-aligned pseudotime trajectories and reveal late-stage heterogeneity. **a)** Violin plots showing transcriptional entropy across pseudotime states in FAPs, MCs, and SCs. Entropy was calculated using ENet-important features (n=228) with the trajectory root selected by the maximum entropy criterion. Black crossbars indicate median values. **b)** Violin plots comparing transcriptional entropy between NSen and Sen cells in scRNA-seq data across cell types. Reduced entropy in Sen cells compared to NSen cells is consistent across all three cell types. P-values from the Wilcoxon rank-sum test. **c)** ROC curves comparing trajectory recovery performance for classifying senescent cells, using four feature sets: variable genes (VARgenes, n=2000), ENet-important genes (ENet, n=228), middle-ranked genes (MIDDLE_228, n=228), and least-important genes (BOTTOM_228, n=228). AUC values are shown for each feature set. ENet genes outperform or match VARgenes across all cell types. **d)** Scatter plots showing correlation between senescence probability predicted by the ENet model (x-axis) and pseudotime from trajectory analysis (y-axis), colored by senescence state. Pearson correlation coefficients and p-values are shown for each cell type. **e)** Boxplots comparing transcriptional entropy among Basal, NSen, and Sen samples in bulk RNA-seq data across three cell types. Individual samples shown as points. Sen samples show reduced entropy compared to NSen, while NSen samples show elevated entropy compared to Basal, suggesting tissue regeneration processes during senescence induction. P-values from Kruskal-Wallis test.

**Extended Data Figure 4:**
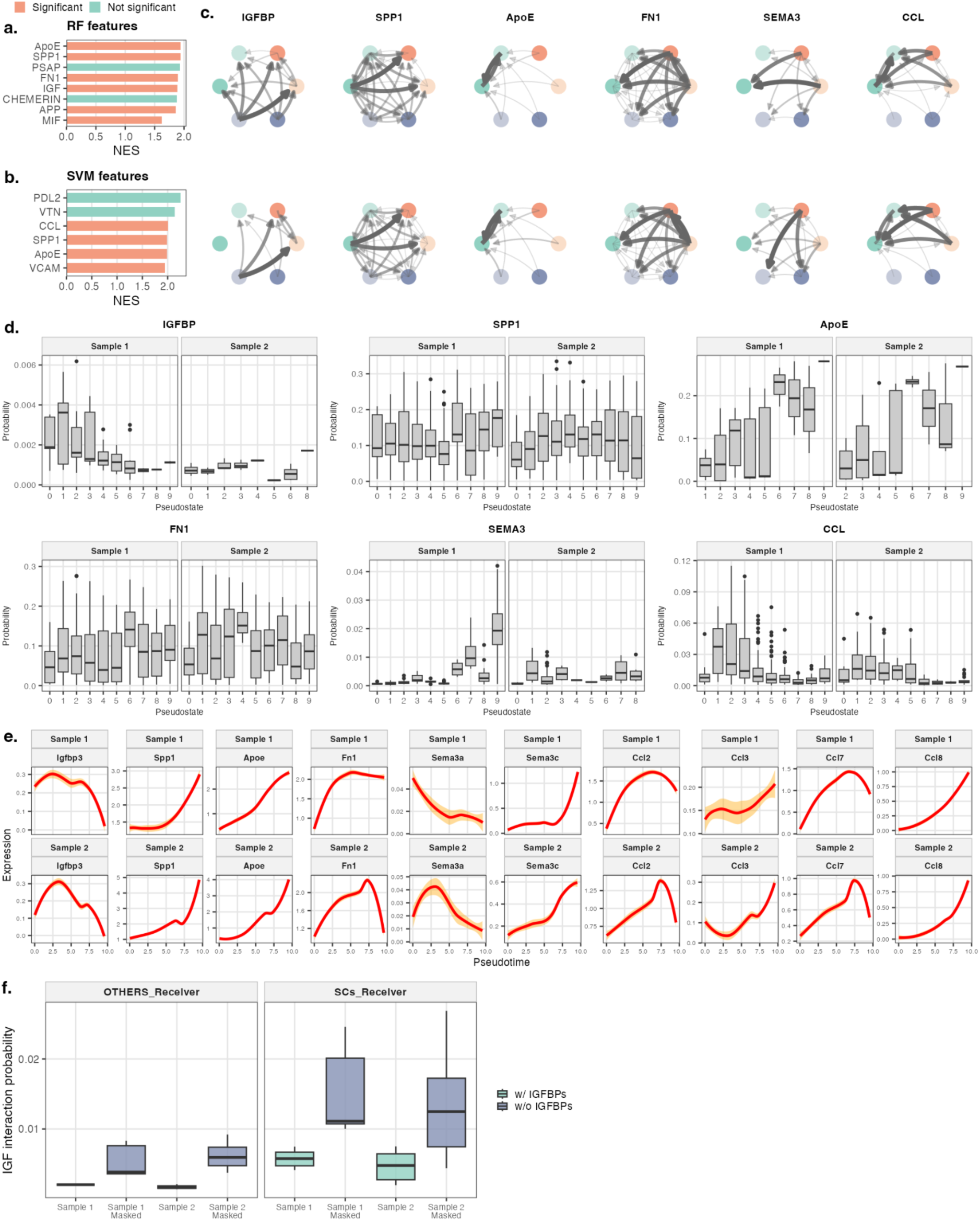
Senescence-associated cell-cell interactions across multiple signaling pathways. **(a)** Barplot showing enriched cell-cell interaction pathways from CellChatDB based on RF-prioritized features, ranked by normalized enrichment score (NES). Significant interactions shown in coral; non-significant in teal. **(b)** Barplot showing enriched interaction pathways based on SVM-prioritized features. Color scheme as in (a). **(c)** Circle plots illustrating cell-cell interaction networks for six ENet-enriched signaling pathways (IGFBP, SPP1, ApoE, FN1, SEMA3, CCL) across senescent (Sen) and non-senescent (NSen) cell types in two biological replicates (top row: Sample 1; bottom row: Sample 2). Nodes represent cell type-senescence combinations colored by identity (FAPs: coral gradient, MCs: teal gradient, SCs: slate blue gradient). Arrow direction indicates sender to receiver; edge width and opacity indicate interaction strength. **(d)** Boxplots showing interaction probability across pseudostates for each pathway in both biological replicates. Interaction probability dynamics along the senescence trajectory reveal pathway-specific temporal patterns. **(e)** Expression trajectories of leading edge genes from ENet-enriched interaction pathways along pseudotime. LOESS-smoothed expression curves (red) with 95% confidence intervals (orange) shown for Igfbp3, Spp1, Apoe, Fn1, Sema3a, Sema3c, Ccl2, Ccl3, Ccl7, and Ccl8 across two samples. **(f)** Boxplots comparing IGF signaling probability with and without IGFBP masking. IGF interaction probability increases when IGFBPs are computationally masked (w/o IGFBPs, slate blue) compared to baseline (w/ IGFBPs, teal), consistent with IGFBP-mediated sequestration and antagonistic modulation of IGF signaling. Left panel shows interactions received by non-SC cell types (OTHERS_Receiver); right panel shows interactions received by satellite cells (SCs_Receiver).

**Extended Data Figure 5:**
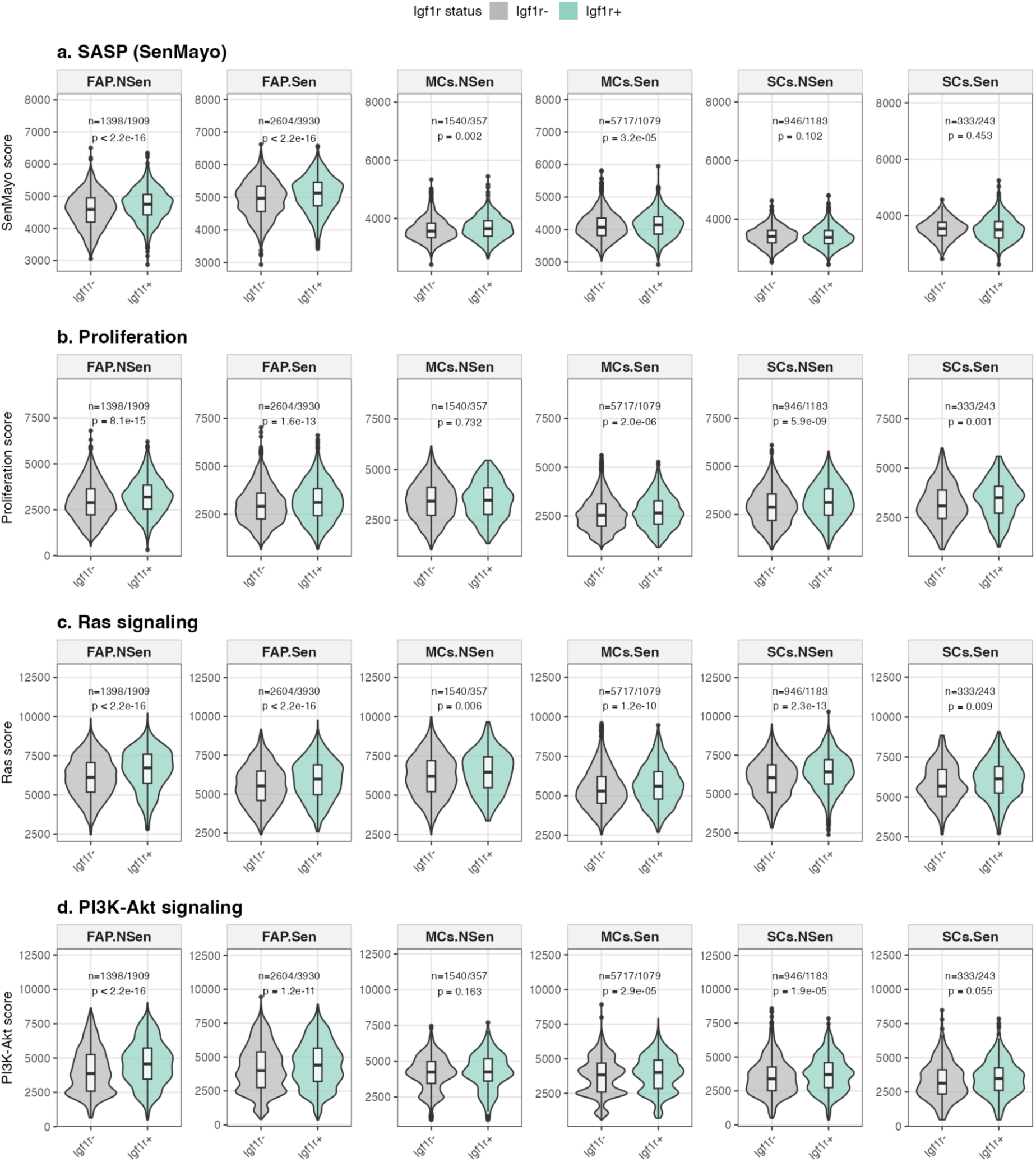
Pathway enrichment scores by Igf1r expression status across cell types and senescence states. Violin plots with embedded boxplots showing the distribution of pathway enrichment scores in Igf1r-negative (gray) and Igf1r-positive (teal) cells, stratified by cell type (FAP, MCs, SCs) and senescence status (NSen, Sen). Data from both biological replicates are combined. Black boxes indicate interquartile range with median line; whiskers extend to 1.5× IQR. **(a)** SASP score (SenMayo gene set). **(b)** Muscle proliferation score. **(c)** Ras signaling pathway enrichment. **(d)** PI3K-Akt signaling pathway enrichment. Statistical comparisons between Igf1r+ and Igf1r-groups were performed using Wilcoxon rank-sum tests; p-values and sample sizes (n = Igf1r-/Igf1r+) are shown for each comparison. Igf1r+ senescent cells show consistently elevated SASP, proliferation, and downstream signaling scores compared to Igf1r-cells, particularly in FAPs and MCs. This figure provides detailed statistical support for the summary heatmap in Figure 4g.

**Extended Data Figure 6.**
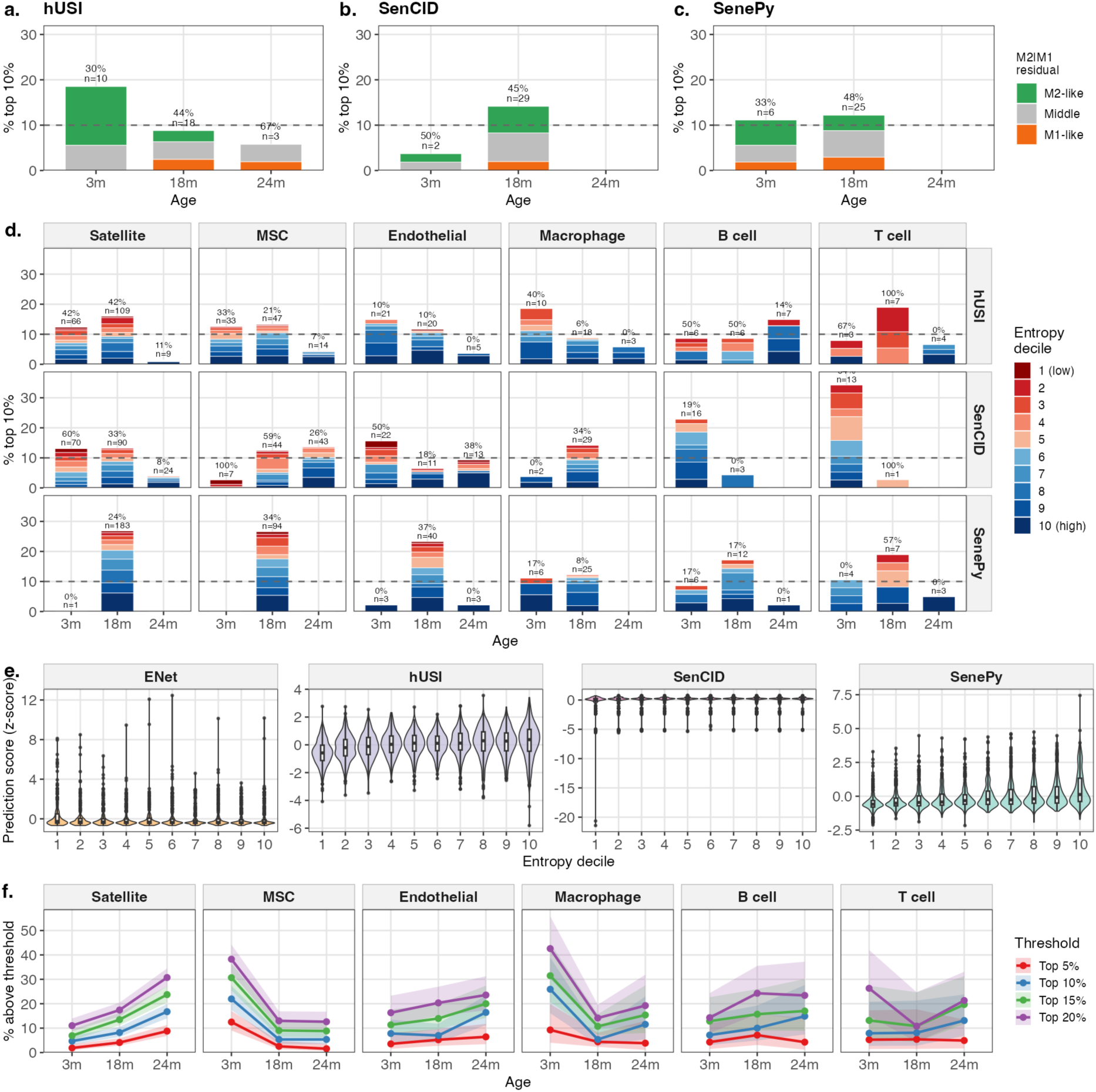
Multi-method comparison of macrophage polarization and entropy-senescence relationships. **(a-c)** Distribution of senescent macrophages by M1/M2 residual classification for alternative senescence prediction methods. (a) hUSI, (b) SenCID, (c) SenePy. Macrophages were classified as M2-like (green), intermediate (gray), or M1-like (orange) based on residuals from M2 versus M1 regression. The age-associated shift from M2-like toward M1-like phenotypes is consistent across all methods. ENet results are shown in Fig. 5h. **(d)** Percentage of cells classified as senescent across entropy deciles, stratified by prediction method (rows) and cell type (columns). Text annotations indicate the percentage of senescent cells in low entropy deciles and the total count. Unlike ENet, the alternative methods do not show consistent low-entropy enrichment in senescent cells. **(e)** Distribution of senescence prediction scores (z-scored within cell type and method) across entropy deciles. Each panel shows one method (ENet, hUSI, SenCID, SenePy). Violin plots with embedded box plots show the distribution of scores at each entropy decile. Lower deciles indicate lower transcriptional entropy. **(f)** Sensitivity analysis of ENet senescence classification at varying thresholds. Each panel shows one cell type; lines indicate the percentage of cells exceeding different thresholds (top 5%, red; top 10%, blue; top 15%, green; top 20%, purple) across ages. Shaded ribbons indicate 95% Wilson score confidence intervals. Age-dependent trends are largely consistent across threshold stringency for each cell type.

## Supplementary Tables

***Supplementary Table S1:*** *Gene sets applied to compute enrichment scores for the classification of senescent versus non-senescent states*.

***Supplementary Table S2:*** *Feature importance scores derived from three machine*-*learning models (Elastic Net, Random Forest, and Support Vector Machine). The table also includes the top 228 features prioritized by the Elastic Net model, selected using the elbow*-*test threshold*.

***Supplementary Table S3:*** *Reduced Gene Ontology Biological Process (GO-BP) enrichment results obtained after semantic similarity filtering, comparing each developmental state to the root state along the pseudotime trajectories in FAPs, MCs, and SCs*.

